# DNA methylation site loss for plasticity-led novel trait genetic fixation

**DOI:** 10.1101/2020.07.09.194738

**Authors:** Takafumi Katsumura, Suguru Sato, Kana Yamashita, Shoji Oda, Takashi Gakuhari, Shodai Tanaka, Kazuko Fujitani, Toshiyuki Nishimaki, Tadashi Imai, Yasutoshi Yoshiura, Hirohiko Takeshima, Yasuyuki Hashiguchi, Yoichi Sekita, Hiroshi Mitani, Motoyuki Ogawa, Hideaki Takeuchi, Hiroki Oota

**Author notes:** A2 Healthcare Corporation, Tokyo, Japan (Kana Yamashita). Stretch store Hataraku, Saitama, Japan (Shodai Tanaka).

## Abstract

Phenotypic plasticity allows organisms to adapt traits in response to environmental changes, yet the molecular basis by which such plastic traits become genetically fixed remains unclear. Here we investigated gut length plasticity in medaka fish (*Oryzias latipes*) through genome-wide methylation profiling, CRISPR/Cas9-mediated deletion, and population genomic analyses. We found that seasonal methylation of CpG sites upstream of the *Plxnb3* is correlated with gut length plasticity, and deletion of this region abolishes plasticity. Additionally, standing variation in *Ppp3r1* is associated with genetically fixed longer gut length in populations lacking plasticity. These results suggest that loss of epigenetic regulation via CpG site reduction triggers the genetic fixation of novel traits. Our findings provide molecular evidence linking epigenetic plasticity and genetic assimilation, advancing understanding of plasticity-led evolution in natural populations.

**One-Sentence Summary:** Seasonal gut-length variation in medaka fish shows how plastic traits evolve into genetic and novel traits.

## Introduction

Genetic and environmental factors that alter phenotypes are key drivers of novel trait evolution. In current evolutionary biology, a novel mutation that contributes to a novel trait spreads in a population through generations if it is advantageous. On the other hand, environmental changes often induce variation in traits, a phenomenon known as phenotypic plasticity (*1*). When such environmentally induced traits are exposed to selective pressures over generations, they may eventually become expressed by subsequent/cryptic genetic mutations, independent of environmental factors, even in natural populations (*2*, *3*). This process, also called the “plasticity-led” evolution (PLE), has been proposed as an alternative pathway of adaptive evolution (*4*). However, whether the existing evolutionary framework can fully explain the PLE process remains unclear (*5*), largely because the molecular mechanisms by which novel traits originating from plasticity become genetically fixed are still unknown.

The PLE process is characterized by the expression of phenotypic plasticity in ancestral lineages and the subsequent genetic fixation of these traits in derived lineages (*4*). In other words, PLE describes how traits initially induced by environmental factors can become genetically fixed, often accompanied by a loss of environmental responsiveness during evolution. To investigate the molecular mechanisms underlying PLE, it is useful to conceptualize the process in two steps: (1) the loss of phenotypic plasticity and (2) the genetic fixation of the induced trait. Phenotypic plasticity is widely considered to arise from changes in gene expression patterns, often mediated by epigenetic modifications such as DNA methylation (*6–8*). The PLE hypothesis further posits that cryptic genetic mutations can fix plastic traits originally induced by environmental change, allowing these traits to be inherited by subsequent generations. If this is the case, genetic mutations responsible for the novel trait may occur within regions of epigenetic modification associated with phenotypic plasticity. To explore the molecular basis of these two steps and their involvement in the PLE process using medaka as a model, we focused on gut length—a trait that exhibits pronounced phenotypic plasticity, is known to enhance nutrient absorption efficiency in response to the feeding environment (*9*), and displays genetic variation across various animal species (*10*, *11*).

The medaka (*Oryzias latipes*), also known as Japanese rice fish, is a small, omnivorous freshwater fish distributed across a wide latitudinal range in East Asia. Local populations of medaka adapt genetically to the various environments of each region, and show geographic phenotypic plasticity and latitudinal variation in traits associated with gut length (*12*, *13*). Moreover, the evolutionary history of medaka is well-studied at the genomic level (*14*, *15*), and various molecular biological techniques can be applied to them (*16*). These features make medaka an excellent model to test whether a focal trait meets the key criteria of PLE (phenotypic plasticity in the ancestral lineage and genetic fixation in the derived lineage) (*2*), allowing us to investigate the molecular mechanisms underlying the PLE process. In this study, we aimed to identify the PLE process underlying their gut length differences and reveal the molecular basis and evolutionary process underlying the transition from plasticity to genetic fixation.

By leveraging medaka’s evolutionary-ecological features, coupled with two genome-wide approaches and subsequent genome editing, allowed us to elucidate the molecular mechanism of PLE underlying gut-length variation in local medaka populations. First, our measurements of gut length revealed both seasonal and geographical variation, demonstrating that the evolution of the medaka gut fulfills the criteria of PLE. Second, genome-wide methylation and population genomic analyses identified candidate genes involved in the two key steps of the PLE process. Finally, functional and molecular evolutionary analyses of these genes showed that a longer gut became genetically expressed through fixation of mutations after the loss of gut-length plasticity associated with a reduction in CpG sites. Although the molecular mechanisms underlying the transition from a plastic trait to a genetically fixed trait have remained largely unknown, our study reveals that the loss of epigenetic modification sites can trigger PLE and expose cryptic genetic variation responsible for novel traits.

## Results

We first verified whether or not the medaka gut shows the phenotypic plasticity on ancestral lineages which are one of the requirements of PLE. We investigated the gut length, collecting wild medaka in three rivers, the Kabe, Ejiri, and Shihodo, in Kagawa Prefecture, Japan (Fig. 1A and table S1). We measured the gut length of 18–23 medakas in each river in two seasons, winter and summer, over a 2-year period, and found statistically significant changes in the gut length relative to standard length (body length) in the Kabe and Ejiri rivers between seasons (fig. S1). Especially in the Kabe river, the medaka in the winter had significantly shorter guts than did those in the summer, which was observed repeatedly over two years (Fig. 1B, Bayesian generalized linear mixed model [GLMM], estimate = -0.44±0.05, 95% credible interval [CI] did not include zero, table S2 and fig. S1). By plotting body and gut lengths in each season, we found no correlation between them in the summer when the gut is longer (fig. S2). To investigate whether or not this change in gut length caused by mixing genetically distinct populations, we analyzed genetic diversity change between winter and summer based on the single nucleotide polymorphisms (SNPs) in the mtDNA and chromosomes and did not detect any noticeable changes of genetic diversity including gene flow from other populations (fig. S3). Taken together, with the lifespan of medaka in the wild (1–1.5 years) (*17*), no overwintering twice (*18*) and no significant difference in the body length between seasons (fig. S1), these results indicated that the gut length changes in the Kabe river medaka must be phenotypic plasticity within individuals in response to the environment.

**Fig. 1.**
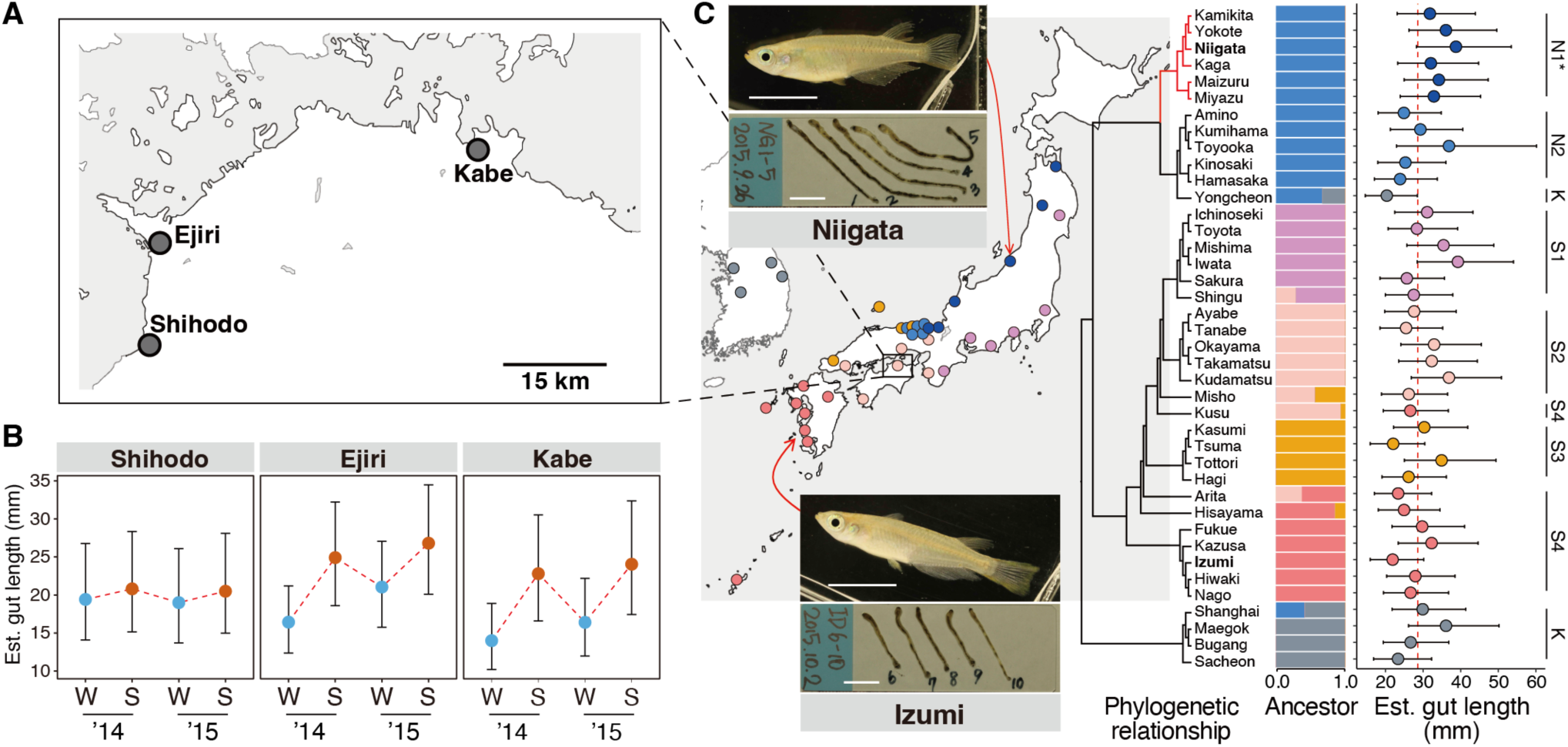
Phenotypic plasticity and genetic variation of the gut length in medaka. (A) A map of sampling locations (gray circles) in Kagawa Prefecture, Japan. (B) Seasonal gut length changes in the three rivers. The blue and orange dots, their error bars indicate the mean gut length and 95% CI (credible interval) after adjusting for the effect of body length in winter (W) and summer (S), respectively, as estimated by Bayesian GLMM. The red dash line joins the mean in each season representing the seasonal variation. (C) Geographic variations of gut length in 40 local populations with example photos (white bars: 1 cm). The phylogenetic tree and ADMIXTURE barplot were constructed using genome-wide single nucleotide polymorphisms (SNPs) published in our prior study (*14*). The right plot shows the geographical variation in gut length corrected for body length using a GLMM across medaka populations. Their points and error bars were fitted values with 95% CIs from the Bayesian GLMM. Abbreviations on the far right: N1, 2, S1–S4, and K are NJPN (Northern Japanese group) 1, 2; SJPN (Southern Japanese group) 1–4; and KOR (Korean/Chinese group), respectively. The asterisk indicates that the NJPN1 medaka (the red branches in the phylogenetic tree) have a significantly longer gut than do the other subgroups.

Gut-length changes are often related to food source (*11*, *19*). To examine whether the food source changes between the seasons, we conducted stable isotope analysis of the caudal fins from the same medaka used for gut-length measurements in three rivers. The stable isotope analysis, based on δ^13^C and δ^15^N, showed no correlation with gut length (fig. S4), indicating that neither food source diversity nor trophic level (i.e., carnivorous habits) strongly influenced gut-length plasticity. To further test whether the food source affects gut length, we transported wild-captured medaka from the Kabe river to our outdoor breeding facility and bred them under artificial conditions. The outdoor breeding experiment showed the same plasticity even under the condition of feeding with artificial food (fig. S5, table S3), indicating no relationship between gut-length plasticity and food source. Therefore, gut-length plasticity may be caused by other environmental factors that we could not measure, such as the amount of food consumed.

We subsequently verified whether or not the medaka gut shows genetic fixation on derived lineages, which is also one of the requirements of PLE, using wild-derived laboratory stocks consisting of 81 geographical local populations. Our previous genome-wide SNP study showed that these stocks cover a genetic diversity of wild medaka and were genetically divided into seven subgroups (SJPN1-4, NJPN1, NJPN2, and KOR) derived from three lineages from the northern Japanese (NJPN), the southern Japanese (SJPN), and the Korean/Chinese (KOR) medaka groups (*14*). Based on the population genetic background of these stocks, we sampled 10 individuals from 40 out of 81 populations to avoid bias from population structure (table S4) and measured the gut length dissected. All local populations were reared in the same environment, including foods, in our outdoor facility. Nevertheless, the NJPN1 subgroup showed a significantly longer gut compared to the average body length than did those of the other subgroups (Fig. 1C, Bayesian GLMM, estimate = -0.11 – -0.26, 95% CIs did not include zero, table S4 and S5), indicating that a longer gut length of the NJPN1 subgroup is fixed by genetic factors common within this subgroup. Meanwhile, the gut lengths within the SJPN and KOR groups vary, indicating that the gut length diversity in the SJPN and KOR groups is defined by genetic factor(s) or environmental factors. As in the wild medaka in Kagawa (Fig. 1B), to examine whether or not the gut of these stocks also showed seasonal plasticity, we compared gut length between seasons using the available stocks from each group (fig. S6A). In NJPN1, the gut length was longer than the body length regardless of the season, although an exception was observed. In NJPN2, there was no significant variation in gut length by season, except in the Toyooka population. In contrast, SJPN, which shares ancestry with the Kagawa medaka, showed large seasonal fluctuations in gut length. Our GLMM analysis showed that the genetic background of SJPN interacted with the seasons and influenced the gut length (fig. S6B, table S6). Together with the seasonal plasticity of the Kagawa medaka which belong to the SJPN group (Fig. 1B), these results indicate the phenotypic plasticity of gut length exists at least in the SJPN lineage. Therefore, based on the evolutionary history of medaka (*14*), the ancestors of medaka likely had phenotypic plasticity in gut length, and the derived subgroup (i.e., NJPN1) of medaka probably acquired a longer gut length to adapt. Thus, we concluded that the evolution of the medaka gut likely meets the requirements for PLE.

Under the PLE hypothesis, DNA methylation, which occurs as environmental responses to seasons and is stable over a long time if the seasonal environment remains unchanged (*20*), is a potential candidate as a molecular basis of seasonal phenotypic plasticity. To test whether the DNA methylation is related to gut length plasticity, methyl-CpG binding domain sequencing (MBD-seq) was used to detect the methylation of DNA extracted from the gut of Kabe river medakas which we observed as exhibiting the strongest seasonal plasticity (Fig. 1B, fig. S1). The MBD-seq analysis revealed hypo- or hypermethylated regions associated with winter or summer, respectively, for 2 years (Fig. 2A and fig. S7). We detected a total of 206 seasonally varying methylated regions: 14 and 192 regions hypermethylated in winter and summer, respectively. We annotated these differentially methylated regions and explored the genes in their vicinity (fig. S8). A total of 71 genes were present in the regions within 2 kb (kilobase) upstream and downstream (two and 69 genes in winter and summer methylated regions, respectively) (Fig. 2A, table S7). We conducted enrichment analyses using the gene lists using Reactome (https://reactome.org/) and Gene Ontology (GO) (http://geneontology.org/) analyses, and found that the semaphorin-plexin signaling related genes (*Plexin a3* [*Plxna3*], *Plexin a4* [*Plxna4*], and *Plexin b3* [*Plxnb3*]) involved in the neuronal axon guidance were significantly enriched by both analyses (Fig. 2A, table S8). Notably, the *Plxnb3* gene, which is involved in the inhibition of neurite outgrowth(*21*), had a CpG island (CGI) in the upstream region showing seasonal methylation (Fig. 2B). We then performed both the whole-genome bisulfite sequence (WGBS) and the targeted bisulfite sequence (tBS) to determine which CpG sites in the upstream regions were seasonally methylated. The WGBS analysis reconfirmed methylation changes in 63 regions detected by the MBD-seq (fig. S9), and the Reactome analysis again showed enrichment of the semaphorin-plexin signaling-related pathway using the list of adjacent genes in these regions (table S7). In the tBS analysis, a statistically significant seasonal difference in methylation was observed at the second CpG site (fig. S10), although a G/A SNP was detected at the 5th CpG site which led to the difference in the methylation level. Furthermore, the methylation of the second and the adjacent CpG sites were significantly correlated and seasonally varied (fig. S11). These correlations were not observed in the populations collected from the Ejiri and Shihodo rivers where seasonal changes in gut length were unclear (fig. S11). A real-time quantitative polymerase chain reaction (PCR) showed that the gene expression of *Plxnb3* in summer was lower than that in winter (*P* < 0.01), indicating the gene expression was correlated with the seasonal methylation on the upstream region of *Plxnb3* (Fig. 2C). Considering the *Plxnb3* gene function and correlation, the upstream region showing the seasonal methylation might be the molecular basis for regulating the gut length plasticity.

**Fig. 2.**
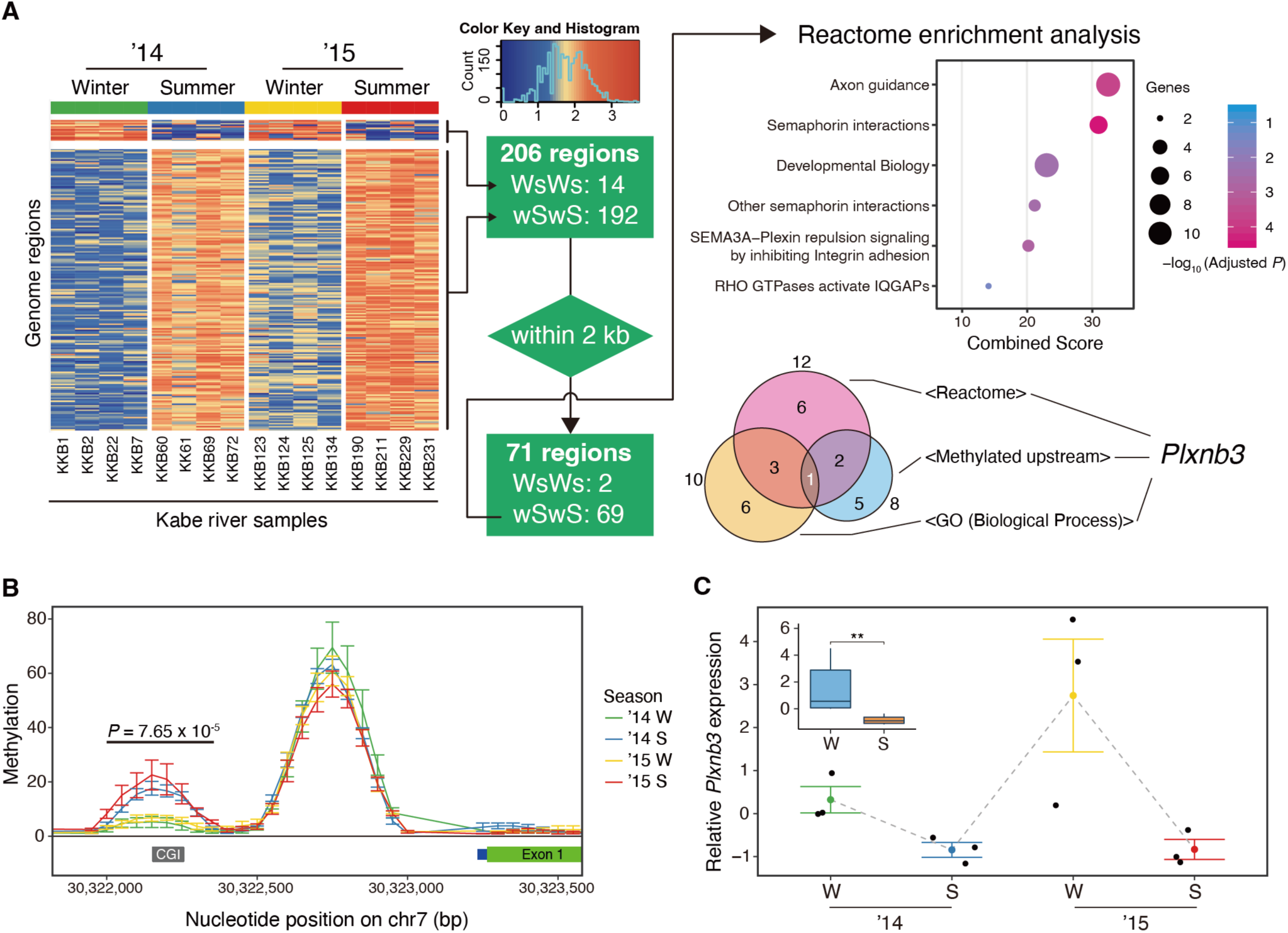
DNA methylation map and *Plxnb3* gene expression pattern correlated with winter and summer. (A) Two hundred six differentially methylated regions (DMRs) were filtered using Reactome, GO analyses, and the position of the methylated region for the gene. The color difference of the heat map shows the difference in the degree of methylation, i.e., orange and blue show the hypermethylated and hypomethylated region, respectively. The four different colors of bar on the top of heatmap represent the combinations of season and year, which are also assigned to Fig. 2B and 2C. WsWs and wSwS represent hypermethylation in winter and summer, respectively. (B) The upstream region of *Plxnb3* showed seasonal methylation, which included the CpG island (denoted by the gray CGI bar). The y-axis displays normalized read counts, plotted as the mean ± SE for 50 bp non-overlapping windows. A significant difference was detected by an analysis of deviance adjusted the Benjamini-Hochberg procedure. (C) *Plxnb3* gene expression was suppressed in the summer (error bars indicate mean ± SE from three biological replicates). The inset graph shows the difference in the gene expression between seasons for two years (2014 + 2015) at the time. A significant difference in gene expression between seasons was detected using the Wilcoxon rank-sum test (\*\**P* < 0.01).

The plasticity observed in the wild population was that the change of the gut length responded to the seasonal environment. Thus, if the environment is constant, the gut length will be constant regardless of body length in individuals that show plasticity. In contrast, in individuals that have lost plasticity, the gut length will increase with body length without responding to the constant environment. To examine whether the upstream sequence of *Plxnb3* is associated with seasonal plasticity of the gut, we generated a genome-edited medaka, *ΔupPlxnb3*, in which the region showing seasonal DNA methylation was deleted using CRISPR/Cas9. We then reared a population of these three genotypes obtained by heterozygous mating in a laboratory constant environment (14/10 light/dark cycle at 28°C) and examined the relationship between body and gut length among the genotypes. At first, testing the segregation ratio of genotypes after a heterozygous mating showed the homozygous and heterozygous rates of medaka with the deletion were about half that compared with the expected segregation rates (Fig. 3A, *P* = 0.011 using the chi-square test). Next, comparing genotypes, no significant difference in gut length was detected, but a significant difference in body length was observed between the wild type and mutant (Fig. 3B, fig. S12). Finally, by plotting body and gut lengths (Fig. 3C, fig. S12) for each genotype, we found dependency between those lengths in the mutants but not in the wild types (Bayesian generalized linear model [GLM], the effect size of the deletion and interaction between standard length and the deletion; estimates = -1.67±0.67 and 0.08±0.03; each 95% CI did not include zero, table S9). These results show that the mutants could have a gut that lengthens in proportion to its body length without responding to the laboratory environment, indicating that the mutants with deletion of the *Plxnb3* upstream sequence lost gut plasticity.

**Fig. 3.**
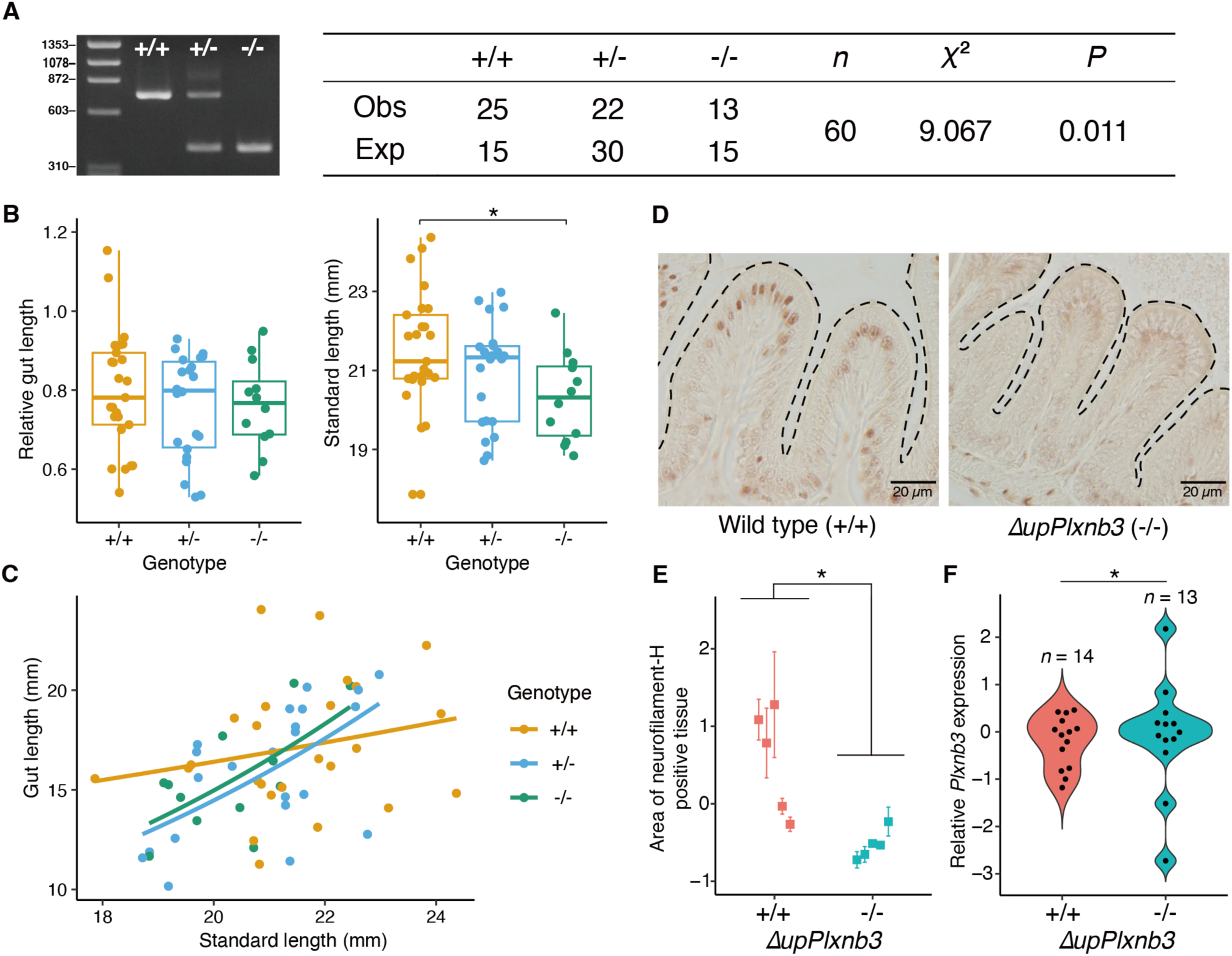
Effects of the deletion of the *Plxnb3* upstream region on gut plasticity. +/+, +/-, and -/-represent the wild type, the heterozygote, and the homozygote of the *Plxnb3* upstream deletant (i.e., the wild type/*ΔupPlxnb3* and *ΔupPlxnb3*/*ΔupPlxnb3*), respectively. (A) The deviation of the segregation ratio among genotypes was evaluated using the chi-squared test. (B) Complete plots of gut length relative to standard length, standard length and (C) their relationship in the heterozygous mating experiment. The genotypes, +/+, +/-, and -/- indicate the wild type/wild type, wild type/*ΔupPlxnb3*, and *ΔupPlxnb3*/*ΔupPlxnb3*, respectively. The asterisk indicates a significant difference between +/+ and -/-genotypes evaluated by using the Steel-Dwass test (\**P* < 0.05). (D) Immunochemical staining revealed the decrease of neurofilament-positive cells at the tip of the intestinal villus. (E) The difference in the degree of immunochemical staining (D) was statistically quantified and detected using the Wilcoxon rank-sum test (\**P* < 0.05). Error bars indicate mean ± SE from 4–6 technical replicates. (F) *Plxnb3* expression showed the uneven variance between the wild types (+/+) and the mutants (-/-), which was evaluated using the F-test (\**P* < 0.05), indicating the unstable *Plxnb3* expression in the mutant gut. The orange and blue in (E) and (F), indicate the wild type and the homozygote of the *Plxnb3* upstream deletant, respectively.

The lower survival rate and shorter body length of the mutants led us to hypothesize that the gut of each individual lacking the upstream sequence of *Plxnb3* undergoes some histological changes. Therefore, we investigated the distribution of neuronal axons involved in the *Plxnb3* by immunostaining, using neurofilament H protein as a marker (Fig. 3D) (*22*). Compared with the wild-type intestinal villi, we found that the immunohistological signal disappeared at the tip of the intestinal villus in the mutants (Fig. 3D and E). Moreover, the *Plxnb3* expression level of the mutants showed significantly greater variance than that of the wild type, suggesting reduced stability in gene expression regulation (Fig. 3F). These results suggest that the seasonal methylated region includes the regulatory region of *Plxnb3*, which that deletion may lead to the disruption of neural mechanisms that senses the seasonal changes of feeding environment (such as sensory-cell mediated food detection (*23*)). Then, it may have caused inadequate nutrient absorption adapted to the environmental change, resulting in a short body length and a lower survival rate (Fig. 3A and B). Overall, our data indicate that the upstream region of *Plxnb3* contributes to the plasticity of gut length, and it is plausible that the seasonal methylation that occurs in the upstream region is linked to the plasticity of gut length.

Under the PLE hypothesis, a genetic mutation should occur and fix a plasticity trait. Because the NJPN1 subgroup exhibited a long genetically fixed gut (Fig. 1C), the causative genetic mutations should be presumed to be highly frequent in the subgroup. In order to examine which gene is responsible for the longer gut of the NJPN1, 62 K SNPs obtained by RAD-seq (*24*) were used to search genomic regions that were differentiated in the NJPN1 subgroup and associated with the gut length (Fig. 4A). Among all candidate loci, we found that the strongest signal associated with gut length was detected in the *Protein phosphatase 3 regulatory subunit B, alpha* (*Ppp3r1*) gene on chromosome 1. This gene is particularly relevant as it is involved in neuronal axon outgrowth (*25*, *26*) and intestinal development (*27*), and showed high genetic differentiation. Bayesian GLMM analysis with population stratification as a random effect confirmed that the T/T genotype had a significant effect on gut length compared to the C/C genotype (Fig. 4B, table S10). This mutation was a synonymous SNP, with no linked non-synonymous SNPs in the coding region within the NJPN1 subgroup (fig. S13), which predicted that the changes in the *Ppp3r1* expression contributed to the gut length. Given that a previous study of conditional *Ppp3r1* KO mice showed hypoplasia of the small intestine (*27*), we could hypothesize that the difference of this gene expression in medaka would lead to geographical differences in gut length. To test this hypothesis, we examined the relationship between *Ppp3r1* expression levels and gut length using outdoor-reared wild-derived laboratory stocks in the winter because a remarkable difference in gut length was expected. We found that the gut length was significantly correlated with the *Ppp3r1* expression level across different genetic backgrounds (Fig. 4C, fig. S14, table S11), and the NJPN1 medakas with a homozygous T allele showed higher *Ppp3r1* expression than other subgroups with a homozygous C allele (Fig. 4C, *P* < 0.01). In addition, in the Iwata and Ichinoseki populations with the same genetic background (SJPN1), a homozygous T allele showed higher *Ppp3r1* expression than a homozygous C allele (fig. S15A, B), and the gut length was significantly correlated with the *Ppp3r1* expression level (fig. S15C). Furthermore, the significant difference in relative gut length between T and C allele homozygotes was confirmed in independent samples from Iwata and Ichinoseki populations (fig. S15D), consistent with the pattern shown in Fig. 1C. These results indicate that the mutation associated with the up-regulation of the *Ppp3r1* expression is involved in the genetic fixation for the longest gut among the medaka subgroups.

**Fig. 4.**
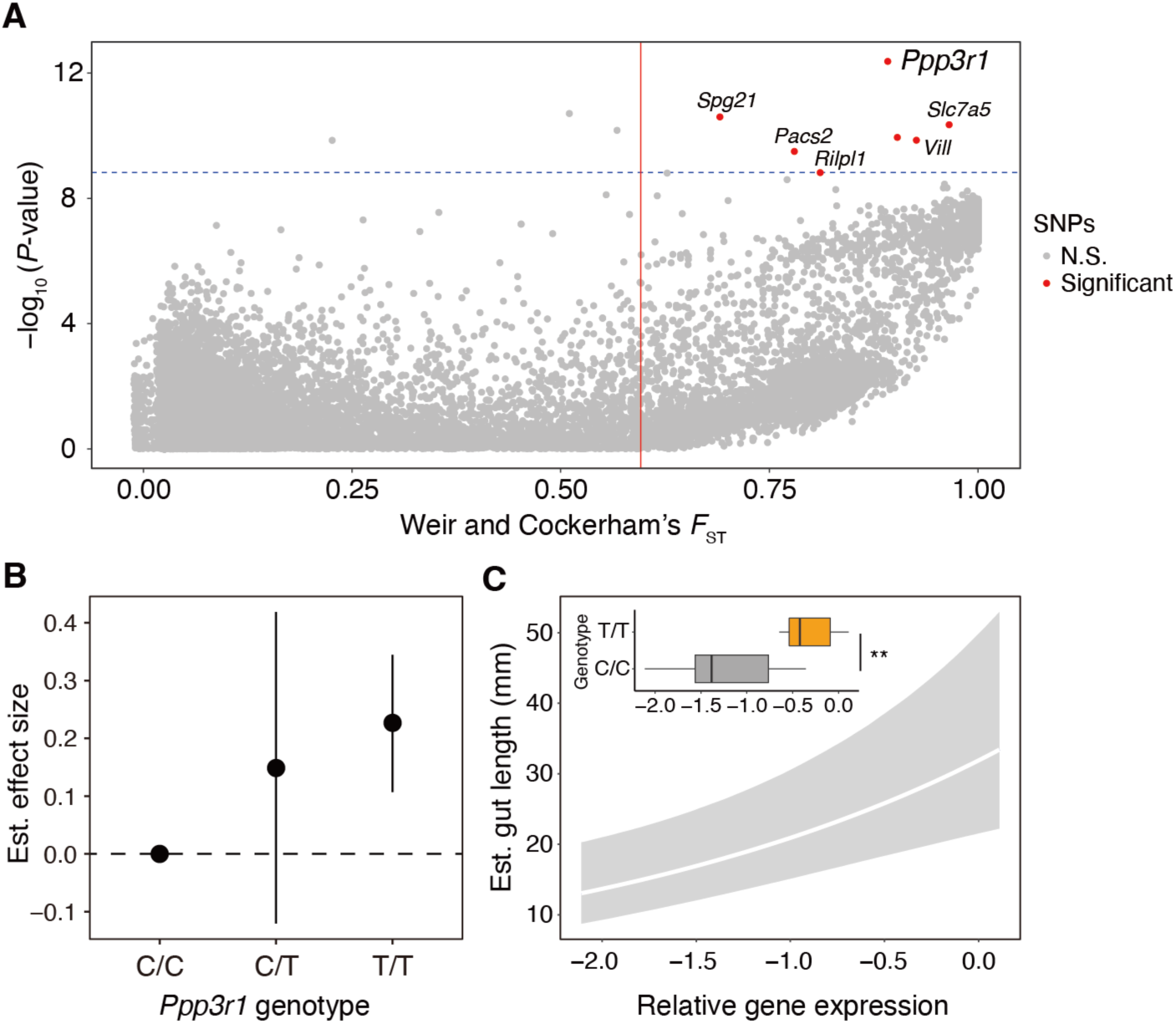
Differentiation and association SNPs with gut length across the medaka populations. (A) *F*_ST_ was calculated between NJPN1 and the others. The red line indicates 0.59592 Weir and Cockerham’s weighted *F*_ST_. Association *P*-values with gut length were calculated using 22,235 SNPs filtered by a minor allele frequency of >0.05. The blue dashed line shows the *P*-value in which the top 10 SNPs distinguished from the others (table S12). The red points show that significant SNPs fulfilled the above criteria. (B) The effect size of *Ppp3r1* genotypes of gut-length associated SNPs (Cytosine [C] or Thymine [T]) was estimated from the posterior mean of a Bayesian GLMM and is given relative to the model intercept at the C/C genotype (dashed line). Their estimates, calculated with population stratification as a random effect, are in table S10. Vertical bars indicate 95% CI. (C) *Ppp3r1* expression was significantly correlated with gut length corrected by standard length, estimated by Bayesian GLMM (effect size of the *Ppp3r1* expression: estimate =0.42±0.12; 95% CIs did not include zero, full data in fig. S14). Line and shaded area are fitted value and 95% CIs from the Bayesian GLMM. The inset graph shows the difference in the *Ppp3r1* expression between the C/C or T/T genotypes, which were examined using winter-sampled individuals. The statistical significance was evaluated using the Wilcoxon rank-sum test (\*\**P* < 0.01).

Finally, we analyzed the molecular evolution of two identified regions based on the medaka population phylogeny. First, we determined the *Plxnb3* upstream sequences to estimate the evolution of methylated regions associated with plasticity in the medaka populations (Fig. 5A, fig. S16). A comparison of the upstream sequences revealed that the number of CpG sites varied between the populations, indicating nucleotide substitutions resulted in the gain and/or loss of CpG sites (fig. S17). Ancestral sequence estimates indicated that the common ancestor of SJPN and NJPN had eight CpG sites in this upstream region, and the number of CpG sites diversified from seven to 10 upon branching to each SJPN lineage. On the other hand, the number of CpG sites in the *Plxnb3* upstream was reduced to six in the common ancestor of NJPN1 and NJPN2, which did not meet the criteria of CpG islands (GC >50%, Obs/Exp >0.6, and length >100 bp). We subsequently examined our RAD-seq data and observed the *Ppp3r1* mutation in the NJPN1 subgroup and the SJPN1 Iwata population, which occur both on the Sea of Japan and Pacific Ocean sides of the eastern Japanese archipelago, respectively. These results suggest that the mutation appeared at least in the common ancestor of SJPN and NJPN that spread throughout the Japanese archipelago and subsequently existed as a standing variation. Considering the phylogenetic relationship among groups (Fig. 1C, fig. S18), the mutation in *Ppp3r1* must have spread in NJPN1 after the number of CpG sites in the *Plxnb3* upstream decreased in the common ancestor of NJPN1 and NJPN2.

**Fig. 5.**
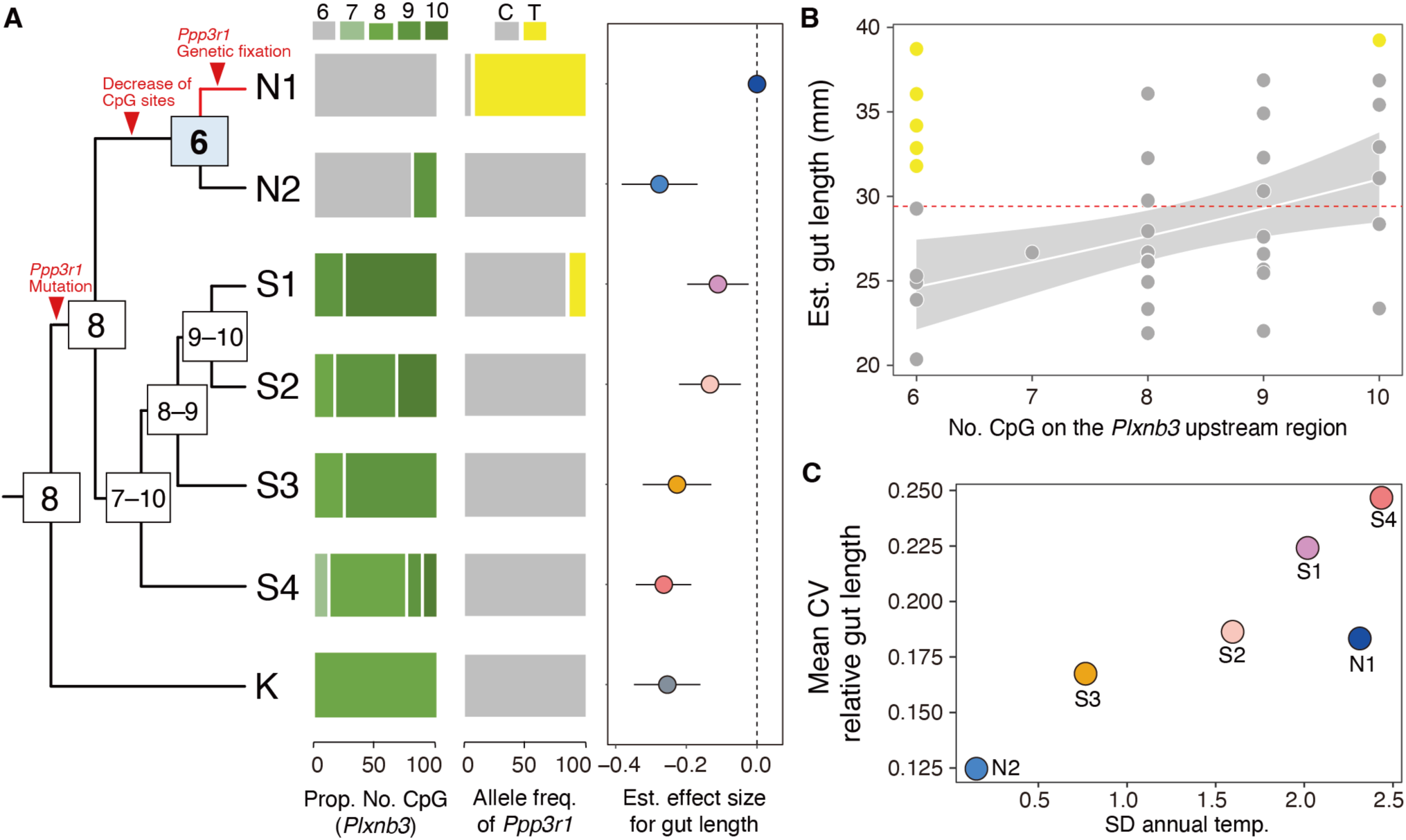
Evolutionary relationship between the *Plxnb3* upstream region and the *Ppp3r1* mutation. (A) The number of CpG sites increased or decreased at each node on the population tree. The red arrowheads indicate evolutionary events, which were inferred from the current proportion of the number of CpG sites on the *Plxnb3* upstream region and the allele frequency of gut-length associated SNPs on the *Ppp3r1* (gray: C-allele; yellow: T-allele). The gradient from gray to dark green at the top of the horizontal bar graph indicates the number of CpG site on the *Plxnb3* upstream region. The right plot shows the effect sizes of each genetic background estimated as a fixed effect in the Bayesian GLMM framework. The effect size is given relative to the model intercept at NJPN1 (dashed line), and each 95% CI of effect size did not include zero, indicating the long gut are fixed genetically in the NJPN1 subgroup. (B) A correlation between the number of CpG sites in the *Plxnb3* upstream and gut length in each local population. The estimated gut length values shown are from the same dataset as Fig. 1C, corrected for body length. Gray and yellow points indicate C- and T-alleles of *Ppp3r1* fixed in each population, respectively. The white line and shaded area are fitted values and 95% CIs from the Bayesian GLMM. (C) A relationship between gut length and annual temperature among subgroups (*r* = 0.869, *P* = 0.025, Pearson’s product-moment correlation test). X-axis and Y-axis indicate the standard deviation (of annual temp.) of their original habitats and the average of coefficient of variation (of relative gut length) within genetic group, respectively. The abbreviations and colored points in (A) and (C) are the same as those in Fig. 1C.

To investigate the relationship between *Plxnb3* and *Ppp3r1*, we estimated the effects of the number of CpG sites in the *Plxnb3* upstream and the presence or absence of the *Ppp3r1* mutation on gut length using GLMM. We found a positive correlation between the number of CpG sites in the *Plxnb3* upstream and population mean gut length estimated by GLMM among the populations with or without the *Ppp3r1* mutation, respectively (Bayesian GLMM, estimate =0.06, 95% CIs did not include zero, table S13), with populations having more CpG sites tending to show longer mean gut lengths (Fig. 5B). In contrast, the *Ppp3r1* mutation had the effect of inducing a long gut regardless of the number of CpG sites in the *Plxnb3* upstream. Even populations with the minimum number of CpG sites displayed long gut lengths if they possessed the *Ppp3r1* T-allele (Fig. 5B), and this mutation effect was statistically significant in the GLMM analysis (table S13). Furthermore, the plot (Fig. 5B) showed that the variability in gut length was small for each *Ppp3r1* genotype when the number of CpG sites was lowest. These results suggest that the number of CpG sites in the *Plxnb3* upstream is involved in the expression of plasticity. Supporting this possibility, we found that NJPN1 and NJPN2 medaka with the lowest CpG sites lost seasonal plasticity in gut length (fig. S6). Furthermore, the *Ppp3r1* mutation was not observed in the Kabe river populations showing seasonal plasticity but in the NJPN1 wild population (fig. S19). These additional results more strongly suggest the gut-length plasticity and genetic fixation of a long gut are associated with the number of CpG sites in the *Plxnb3* upstream and *Ppp3r1* mutation, respectively. Taken together with the molecular evolution of these two regions, our data are consistent with the possibility that the mutation spread to the NJPN1 subgroup after loss of plasticity (called “genetic assimilation” (*28*)) due to reduction of the number of CpG sites in the *Plxnb3* upstream, after which a long gut became expressed in northern populations due to the standing variation of *Ppp3r1* (Fig. 5A).

## Discussion

Epigenetic change enables a single genotype to promote and maintain the phenotypic variation. In this study, we hypothesized that DNA methylation regulates phenotypic plasticity and that loss of the CpG site to be methylated leads to loss of phenotypic plasticity, and found the correlation between seasonal variation in gut length and methylation of the upstream region of the *Plxnb3* gene in medaka (Fig. 2). We also found that deletion of the upstream region of *Plxnb3* leads to a loss of gut plasticity (Fig. 3). Furthermore, a seasonal variation in gut length was not observed in populations with the lowest number of CpG sites (fig. S6). These results indicate that the seasonal variation of CpG site methylation in the upstream region of *Plxnb3* contributes to the gut-length plasticity in medaka. In addition, it is not because substitutions of the CpG sites, that were seasonally methylated, resulted in the loss of gut length plasticity, since those CpG sites are conserved among medaka populations (fig. S17). Thus, the loss of plasticity may not be due to the loss of CpG sites with seasonal methylation but rather to the loss of nearby CpG sites that are stably methylated through seasons. Considering that methylation of promoter regions affects transcription factor binding (*29*), the decrease of CpG sites in the upstream region of *Plxnb3* would weaken the influence of seasonal CpG methylation on transcription factor binding. In other words, constantly methylated CpG sites in the upstream region may have limited the binding of transcription factors only to seasonally methylated CpG sites, consequently making the effects of seasonal methylation of CpG sites more obvious. Under this hypothesis, it can be predicted that the difference in the number of stably methylated CpG sites in the upstream region results in the variability in the *Plxnb3* gene expression. In practice, the variability of the *Plxnb3* gene expression in the population with more CpG sites was smaller than that in the population with fewer CpG sites (fig. S20). Although it remains to be addressed whether or not the stably methylated CpG sites enhance the influence of the seasonal methylation of CpG sites in the vicinity, our results provide strong evidence that DNA methylation is involved in regulating plasticity and that loss of CpG sites results in the loss of plasticity.

Genome-wide SNP analysis indicated that a *Ppp3r1* mutation, which is present as a standing variation, contributed to genetically fixing of the long gut observed in NJPN1 (Fig. 4). This gene is the most crucial candidate because it is involved in neuronal axon outgrowths like *Plxnb3* and small intestine formation (*27*). Because increase in the *Ppp3r1* gene expression strongly correlates with gut elongation (Fig. 4 and fig. S15), a mutation responsible for gene expression might exist in the regulatory region, which is linked to the synonymous mutation in *Ppp3r1* found in this study. Moreover, NJPN1 showed the *Ppp3r1* expression higher than the other subgroups (fig. S15A), suggesting that lineage-specific mutation(s) may be present. On the other hand, this associated mutation (or a linked unknown mutation) explains only the gut length variation in NJPN1 and a part of the SJPN1 subgroup (Fig. 5). The long guts in the other subgroups may be attributable to genetic variations originated by other mutations that occurred and/or were selected independently in each population or a plastic variation induced by environmental pressures under artificial rearing conditions. To address this matter, crossbred experiments will be necessary in future studies. Nevertheless, it is an important finding to understand the process of PLE that a pre-existing genetic mutation in the genomic region completely different from those involved in gut length plasticity is involved in the genetic fixation of the long gut in medaka.

This study shows that two genes involved in neuronal axons development contribute to the phenotypic plasticity and the genetic polymorphism of gut length in medaka. It was clarified at the molecular level that the plasticity was associated with seasonal methylation of CpG in the *Plxnb3* upstream and that the genetic mutation in *Ppp3r1* spread to populations in which the methylation site had reduced by the mutation (i.e., nucleotide substitution). In short, these molecular mechanisms and evolutionary processes indicate that the mutation-driven evolution occurred in gut length. This differs from “plasticity-driven evolution”, which is based on phenotypic plasticity and in which the adaptive trait is led by natural selection and the following mutation fixes it (summarized as, “selection-driven evolution” in Nei [2013] (*30*)). Furthermore, our results indicate that genetic fixation of a plasticity-led trait would occur under the following process. The phenotypic plasticity regulated by CpG methylation plays the role of a buffer for environmental pressure, which contributes to the accumulation and maintenance of *de novo* mutations as standing variations. Once plasticity is lost, the adaptive mutation is selected from standing variations and then the adaptive phenotype is genetically fixed. Our data indicate that this process is likely to be the PLE observed in nature (not regarding a polyphenism (*31*), which is one of the outcomes of plasticity-led evolution (*32*)), and the hereditary phenomenon of the environmentally induced trait could be explained by the genetic framework of loss of CpG sites and fixation with standing variations.

Why has a genetic fixation of the long gut occurred in the NJPN1 medaka? SJPN and NJPN medaka originate in northern Kyushu and the Tajima-Tango region, respectively (*14*). In SJPN medaka, populations with similar genetic backgrounds inhabit the different climatic environments. Because the foraging environment is susceptible to annual temperature changes and the gut length has become varied in response to the average temperature variation (Fig. 5C), the gut length would have been plastically adjusted for efficient digestion and absorption in various foraging environments. On the other hand, NJPN1 is a group that has spread rapidly to the north from small habitats where the environment is relatively uniform (*14*). In the habitat of NJPN2, both temperature and gut length show minimal variability (Fig. 5C), while the NJPN1 habitats show a large temperature variation and short growing period because of the high latitudes of their habitats (*12*). Therefore, the NJPN2 medaka did not need to change their gut length, whereas the NJPN1 medaka must feed during the short summer and prepare for the winter. Under these circumstances for the NJPN1 medaka, maintaining a long gut throughout the seasons may have served as food storage, rather than regulating gut length to increase absorption efficiency as in SJPN medaka. Moreover, the foraging behavior of NJPN1 medaka that has expanded into the higher latitude regions is more frequent than that of medaka in the lower latitude regions (*12*). This suggests that genetic mutations which can change gut length may be also driving the geographic differences in the behavior of foraging. Indeed, the neurofilament protein involved in sensory neural regulation in the gut (*33*) was not stably expressed in medaka that had lost plasticity (Fig. 3D). This result suggests that the gut could not be detecting the appropriate amount of feeding, which may lead to excessive feeding behavior. I.e., the gut-brain interaction could have enhanced the genetic fixation of advantageous mutation.

The identified molecular mechanisms and the above evolutionary inference suggest that plasticity may be lost under a stable environment, and that after loss of plasticity, a favorable mutation can be fixed on foraying into harsh habitats. This phenomenon may appear as though the evolution of acquired traits has occurred in a macroscopic-type view, because it occurs continuously.

## Materials and Methods

### Ethics statement

The Institutional Animal Care and Use Committee of Kitasato University and The University of Tokyo approved all the experimental procedures (No. 4111 and C-09-01, respectively).

### Sampling and measurement of gut length

To detect the seasonal plasticity of gut length, we collected 228 medaka individuals from three rivers in Kagawa prefecture from January 2014 to August 2015 (Fig. 1A). The details are in supplementary table S1. To explore the genetically-fixed gut length in medaka populations, we sampled 400 individuals from 40 wild-derived laboratory stocks that originated from wild populations and consisted of seven genetic backgrounds (SJPN1-4, NJPN1, NJPN2, and KOR) (*14*). Each stock is a descendant of populations collected from each wild habitat in 1983 (*34*) and has been maintained by taking offspring every 1-2 years, close to the generation time in the field (*17*) in independent tanks in common environments (i.e., the same food and feeding times) at our outdoor breeding facility at The University of Tokyo (*14*, *35*) (see fig. S9 in Katsumura *et al.* [2014] (*35*)). The sampled stocks were: SJPN1 (PO.NEH): Ichinoseki, Toyota, Mishima, Iwata, Sakura, Shingu; SJPN2 (Sanyo/Shikoku/Kinki): Ayabe, Tanabe, Okayama, Takamatsu, Kudamatsu, Misho; SJPN3 (San-in): Kasumi, Tsuma, Tottori, Hagi; SJPN4 (Northern and Southern Kyushu): Kusu, Arita, Hisayama, Fukue, Kazusa, Izumi, Hiwaki, Nago; NJPN1 (derived NJPN): Kamikita, Yokote, Niigata, Kaga, Maizuru, Miyazu; NJPN2 (ancestral NJPN): Amino, Kumihama, Toyooka, Kinosaki, Hamasaka; KOR (KOR/CHN): Yongcheon, Maegok, Bugang, Sacheon, Shanghai. The subgroup names in parentheses are defined in our prior study (*14*).

We first took a photo of the whole medaka body lying on the glass slide to measure the standard length (body length) from the most anterior part of the head to the end of the vertebral column. Next, we isolated the gut (from the esophagus to the anus), fixed each one in 4% paraformaldehyde for 1 hour on ice and took photos of the gut laid out on glass slides to measure the gut length as the distance from the esophagus to the anus. About wild medaka samples from three rivers, the caudal fin was dissected for the stable isotope analysis, which was outsourced to Shoko Science Co., Ltd. in Japan. Unfortunately, for 400 samples from 40 wild-derived laboratory stocks, 32 medaka individuals died before this analysis, and 27 individuals were removed because of presenting with an abnormal form due to aging (e.g., spondylosis). Finally, we obtained the standard and gut lengths from photos of 341 individuals (table S4) using Image J software (*36*). Statistical analyses were performed using R software (*37*).

### Detecting gut length changes using a Bayesian GLMM framework

The relationships between gut length and seasons, genetic background, *Plxnb3* upstream, *Ppp3r1* genotypes, and gene expressions were modeled using a GLMM (Generalized Linear Mixed Model) framework with the MCMC (Markov chain Monte Carlo) method in the R package brms (*38*) in R v3.5.2. The gut length was modeled with a gamma error structure and log link function. Fixed effects included the standard length and sex in each analysis, and the random effect was subject identity and genetic background. The MCMC conditions are described in each supplementary table.

### Detecting seasonal methylated regions and filtering using Reactome and GO databases

We extracted genomic DNA from two male and two female guts per season (total 16 guts) from the Kabe river medaka using the phenol/chloroform method described previously (*14*). The quality and quantity of genomic DNA were assessed using 0.5% agarose electrophoresis, a nanophotometer (IMPLEN), and the Qubit BR dsDNA Assay Kit (Thermo Fisher Scientific). After adjusting the DNA concentration to 40 ng/µl, 1.2 µg of DNA was sheared to target 150–200 bp by using two protocols of 400 bp and 200 bp implemented in the S220 Focused-ultrasonicator (Covaris). Sheared DNAs were purified using the MinElute PCR Purification Kit (Qiagen) according to the manufacture’s protocol, and their quality and quantity were measured by the 4200 TapeStation system (Agilent Technologies) and Qubit dsDNA BR Assay Kit (Thermo Fisher Scientific). A total of 500 ng of sheared DNAs was input for methyl-CpG binding domain (MBD) enrichment using the EpiXplore Methylated DNA Enrichment Kit (Clontech) according to the manufacturer’s instruction. Subsequently, 30 ng of methylated DNA was used to generate the NGS (Next-Generation Sequencing) library by using the NEBNext ULTRA DNA Library Prep Kit for Illumina (New England Biolabs) and the NEBNext Multiplex Oligos for Illumina (New England Biolabs) according to the manufacturers’ instructions. The quality and quantity of genomic DNA were assessed using the TapeStation D1000HS screen tape (Agilent Technologies) and the Qubit dsDNA HS Assay Kit (Thermo Fisher Scientific). The sequencing process was outsourced to Macrogen Japan in Kyoto and was performed by Illumina Hiseq 2500 with 51 bp single-end reads to obtain an average of 40 million reads per sample. The data have been submitted to the DDBJ Sequence Read Archive (DRA) database under accession number: DRA010581.

Our single-end reads were filtered using the FASTQ Quality Trimmer and Filter in FASTX-Toolkit ver.0.0.13 (http://hannonlab.cshl.edu/fastx_toolkit/download.html) using the following options: “-t 20 -l 30 -Q 33” and “-Q 33 -v -z -q 20 -p 80”, respectively. The draft genome of the medaka sequenced by the PacBio sequencer (Medaka-Hd-rR-pacbio_version2.2.4.fasta; http://utgenome.org/medaka_v2/#!Assembly.md) was used to align the reads using BWA (Burrows-Wheeler Aligner) backtrack 0.7.15-r1140 (*39*) using the “-n 0.06 -k 3” option. After the mapping process, the multi-mapped reads were removed using SAMtools v1.9 (*40*) and the “-q 1 -F 4 -F 256 -F 2048” option.

The detection of differentially methylated regions (DMRs) was performed using MethylAction (*41*) with “fragsize=200, winsize=50” on R ver.3.4.4 in RStudio ver.1.0.136. First, significant DMRs in one of the groups at least (Adjusted analysis of deviation [ANODEV] *P*-values [Benjamini-Hochberg] <0.05) were explored by comparing between the four groups (14W and 14S, 15W and 15S). Next, the regions that were TRUE in the “frequent column”, which the “frequent” state corresponds to those where the methylation status of the samples is two-thirds or more agreement in each group, were filtered. And then, we extracted regions with patterns that hypermethylation and hypomethylation coincided with seasonal changes repeatedly, as seasonal varying methylated regions. To determine if type I error rates is acceptable, we calculated a false discovery rate (FDR) for each pattern of DMR among the groups by the MethylAction-implemented bootstrapping approach (2500 reps). We then annotated the DMRs (length distribution, overlap with open chromatin location information from ATAC-seq data (*42*), and TE search using RepeatMasker (*43*)) using BEDTools (*44*) and SeqMonk ver.1.48.0 (https://www.bioinformatics.babraham.ac.uk/projects/seqmonk/), and finally listed genes within 2,000 bp upstream and downstream of the DMRs.

To identify the biological processes of genes uniquely associated with each seasonally-correlated region, we performed enrichment analyses based on Reactome pathway, GO terms, and human genome annotations using Enrichr program (*45*, *46*), without correcting for gene size variation. Before this Enrichr analysis, we estimated medaka orthologous genes to human genes using biomart in Ensembl database and ORTHOSCOPE (*47*) because we used an information of gene function in human. Significant Reactome pathways (Adjusted *P*-value <0.05) and the top 10 GO terms for “biological processes” were used as the filter to list the candidate genes. Adding the gene list with the seasonally methylated region on their upstream, we visualized as a Venn diagram and identified the functional gene, *Plxnb3*, associated with seasonal plasticity (Fig. 2A). Finally, we identified a CpG island on upstream of this gene using MethPrimer (*48*) with the following default setting: GC >50%, O/E >0.6, and length >100 bp (Fig. 2B).

### Analyses of whole genome bisulfite sequencing and targeted bisulfite sequencing

We used the same DNA as that used in MBD-seq. The whole genome bisulfite sequencing (WGBS) was outsourced to Macrogen Japan Inc. In brief, genomic DNA was bisulfite converted using the EZ DNA Methylation-Gold Kit (ZYMO RESEARCH). Sequencing libraries were prepared using the Accel-NGS™ Methyl-Seq DNA Library Kit (Swift BIOSCIENCE). Each library was sequenced using NovaSeq6000 (Illumina) to obtain raw data of more than 45 Gb paired-end (about 56 times more than medaka genome).

WGBS analysis was performed using programs: Bismark ver.0.23.0 (*49*) and SeqMonk ver.1.48.0 (https://www.bioinformatics.babraham.ac.uk/projects/seqmonk/). The raw fastq reads (average 51.5 Gb/sample) were filtered using fastp (*50*) with the following option: “-q 20 -n 10 -f 15 -F 15 -t 15 -T 15 -l 20”. The clean fastq reads (average 39.7 Gb/sample) were aligned to medaka reference genome (Medaka-Hd-rR-pacbio_version2.2.4.fasta) with Bismark using the following parameters: “--score_min L,0,-0.6”. The duplicate reads were removed using “deduplicate_bismark” in Bismark. The methylation calls were performed using bismark_methylation_extractor and bismark2bedGraph command in Bismark. The methylation levels were determined for each CpG site within the regions detected by MBD-seq with SeqMonk using the “Difference quantification” function (as a percent of methylated CpGs calls over total calls for each site) and retrieved using the “Annotated probe report” function. Logistic regression filter on probes in MBD-seq detected peaks where summer vs winter had a significance below 0.1 after Benjamimi and Hochberg correction with a minimum number of observations of 10 in all samples. Also, the differences in methylation between season was visualized by SeqMonk.

For targeted bisulfite sequencing (tBS) analysis, we extracted genomic DNA from two male and two female guts per season (total 16 guts) from the Ejiri and Shihodo rivers as well as above. The genomic DNA was bisulfite converted and sequenced using MiSeq (Illumina) as described previously (*51*, *52*). The primer pairs used to amplify the target were: oPlexnb3_MR_Nxt_1F: TCGTCGGCAGCGTCAGATGTGTATAAGAGACAGTATATGTGATGTTTAGAGGATTGG; oPlexnb3_MR_Nxt_1R: GTCTCGTGGGCTCGGAGATGTGTATAAGAGACAGTTTTTAAACTAACTCAAAATTCTAACC. The methylation level for each CpG site was calculated by QUMA (*53*) using the automated shell scripts in Sekita et al. (*51*, *52*) and the following condition to exclude low quality sequences: “Upper limit of unconverted CpHs, 5; Lower limit of percent converted CpHs, 0.95; Upper limit of alignment mismatches, 20; Lower limit of percent identity, 0.95.” Statistical difference in methylation between seasons on each CpG site was evaluated by GLMM using “lmerTest” R package. The methylation (the number of methylated and unmethylated CpG) was modeled with a binomial error structure and logit link function. Fixed effect included the season (winter or summer), and the random effect was sampling year. The short read data of tBS and WGBS have been submitted to the DRA database under the accession numbers: DRA015161 and DRA015162, respectively.

### Finding the high-differentiated and gut-length-associated SNPs

For 341 gut-length-measured medaka, genomic DNAs were extracted from muscle using NucleoSpin Tissue (Macherey-Nagel) according to the manufacturer’s protocol. After the quality check using a Nanophotometer (IMPLEN) and 0.5% agarose gel electrophoresis, six individuals were removed because of low DNA concentrations. For 335 individuals, we arranged the original RAD-seq protocol (*24*) adding an indexing PCR step to adjust the sample size and generated 14 RAD-seq libraries (24 individuals per library, 23 individuals in the last library) by the following method. The designed P1-adaptor included 24 in-line barcodes which had two nucleotide differences in each, and the P1-adaptors and Sbf I-HF (New England Biolabs)-digested DNAs for 90 min at 37°C were ligated by T4 ligase (New England Biolabs). After DNA pooling, DNAs were sonicated using an S220 Focused-ultrasonicator (Covaris) to target 300 bp and purified using the GeneRead Size Selection Kit (Qiagen) according to the manufacturers’ protocols. The End-repair, A-tailing, and P2 adaptor ligation steps were performed using the NEBNext Ultra DNA Library Prep Kit for Illumina (New England Biolabs). After size selection using AMPure XP beads (Beckman Coulter), indexing PCR was performed using Q5 (New England Biolabs) under the following conditions: initial denaturing step at 90°C for 30 sec, 14 cycles of denaturation at 98°C for 10 sec, annealing at 68°C for 30 sec, extension at 72°C for 20 sec, and a final extension step at 72°C for 5 min. The PCR products were purified with AMPure XP beads and then were validated using the Qubit dsDNA HS Assay Kit (Thermo Fisher Scientific) for DNA concentration, 4200 TapeStation (Agilent Technologies) for fragment length, and Miseq (Illumina) for library quality. Finally, RAD-seq data were generated using three lanes of Hiseq 2500 (Illumina) with 51 bp single-end reads settings conducted by Macrogen Japan. The data have been submitted to the DRA database under accession number: DRA010605.

Our single-end reads were filtered by Cutadapt ver1.12 (*54*) using the following options: “-m 50 -e 0.2,” and were demultiplexed by process_radtags (v1.44) implemented in Stacks (*55*) using the following options: “-c -r -t 44 -q -s 0 --barcode_dist_1 2.” After quality filtering and check, one individual (from the Kazusa population) was removed because of low quality reads. The draft genome of the medaka sequenced by the PacBio sequencer (Medaka-Hd-rR-pacbio_version2.2.4.fasta; http://utgenome.org/medaka_v2/#!Assembly.md) was used to align the reads using BWA backtrack 0.7.15-r1140 (*39*) using the “-n 0.06 -k 3” option. After the mapping process, the multi-mapped reads were removed using SAMtools v1.9 (*40*) and the “-q 1 -F 4 -F 256 -F 2048” option. SNP call was performed by Stacks pipeline: pstacks -m 2 --model_type snp --alpha 0.05; cstacks -g -n 1; sstacks -g; rxstacks --lnl_filter --lnl_lim -10 --conf_filter --conf_lim 0.75 --prune_haplo --model_type bounded --bound_low 0 --bound_high 0.1; cstacks -g -n 1; sstacks -g; populations -r 0.5 -p 7 -m 6 -f p_value -a 0.0 --p_value_cutoff 0.1 --lnl_lim -10. Assigned medaka populations into seven genetic groups based on our prior study (*14*), genotype imputation was performed by Beagle (v4.1) using GT format (*56*). Finally, data set was generated using VCFtools (*57*) to merge the seven genotype-imputed data, and included 63,265 SNPs.

To detect high-differentiated and gut-length-associated genes, we calculated Weir and Cockerham’s *F*_ST_ and the degree of association with the gut length using VCFtools (*57*) and PLINK v1.90b3.46 (*58*) (https://www.cog-genomics.org/plink/1.9/) using the SNP data. Although the gut length was highly correlated with the population structure of medaka (Fig. 1C), we did not control the structure information because this reduced the power of the analysis and could lead to false negatives (*59*). To avoid the false positives without controlling the population structure as much as possible, we designed the sampling strategy to choose multiple subgroups from each group and to use balancing samples across subgroup populations to homogenize allele frequencies (*59*). Focusing on SNPs with a minor allele frequency of more than 0.05, we extracted the top 10 SNPs above the global *F*_ST_ value (weighted *F*_ST_) and with the highest -log_10_ *P*-value from 22,235 SNPs. Association of gut length and SNPs were assessed by permutation test with 10,000,000 times resampling, implemented in PLINK v1.90b3.46 (*58*).

### Phylogenetic tree reconstruction using RAD-seq data

We used 22,235 SNPs data, which is the same data above association analysis, for reconstruction of the phylogenetic tree. The data was analyzed via the IQ-TREE program (*60*) to reconstruct a maximum likelihood tree with general time-reversible (GTR) + ascertainment bias correction model and 1,000 ultrafast-bootstrappings (*61*). Then, we used the FigTree ver.1.4.4 program (http://tree.bio.ed.ac.uk/software/figtree/) to visualize the phylogenetic tree.

### Generating a genome-editing medaka by the CRISPR/Cas9 system

The d-rR/Tokyo medaka line was used to generate mutants deleted from the seasonal methylation region. Fish were maintained at 28°C under a 14-h light/10-h dark cycle. We collected the d-rR eggs every morning. Microinjection was carried out in medaka embryos at the one-cell stage using FemtoJet (Eppendorf) and InjectMan NI 2 (Eppendorf). Cas9 protein (500 ng/μL), tracrRNA (200 ng/μL), two crRNAs (each 200 ng/μL), and saturated phenol red solution were mixed in a 4:2:2:1 ratio, and injected into the embryos. The crRNAs were designed to delete the seasonal methylation region of 334 bp using CRISPRdirect (*62*), which identified the specific regions on the medaka genome, and their crRNA sequences were: oPlxnB3upCGI_F1, GAUGCUGUUGCCUGGCCAUAguuuuagagcuaugcuguuuug; oPlxnB3upCGI_R1, GAUGACACAGUGGAGCAUCGguuuuagagcuaugcuguuuug. The crRNAs, tracrRNA, and recombinant Cas9 protein were obtained from FASMAC. After injection, the embryos (F0) were maintained at 28°C using an air incubator. After hatching and maturing, the injected medaka were intercrossed for the identification of the germline founders with mutations in each target locus by genotyping their fertile eggs. The genotyping was performed using PCR and 1.5% agarose gel electrophoresis using EX Taq Hot Start Version (TaKaRa Bio) according to the manufacturer’s protocol and two designed primers: oPlxnb3CpG1F2, 5’-TTGCATCTGGCTTTGCATGAATC-3’; oPlxnb3CpG1R2, 5’-ATAACATCACCATGGCAACAACG-3’. The germline founders (F1) were outcrossed and generated their off-springs (F2). Heterozygous mutation carriers in the F2 generation were identified by above PCR-based genotyping using their fin-extracted DNAs. Finally, sixty F3 individuals generated by heterozygous mating of the heterozygous mutation carriers in the F2 were used for subsequent analyses.

### Analysis of *Plxnb3* upstream-deleted medaka

To examine a phenotype of *Plxnb3* upstream-deleted (*ΔupPlxnb3*) medaka, we conducted three tests: segregation of genotypes, a comparison of gut and body length, and an immunohistological comparison. For these tests, we performed heterozygous mating using the F2 generation and obtained their fertile eggs. We bred the medaka larvae in a tank (W52×D21×H23cm) and raised them until 2 weeks after hatching, divided the juveniles into groups of four to six individuals, and transferred them to 2 L tanks to raise them for an average of 103 days after hatching. Each genotype was determined using the PCR-based method described above, and the segregation ratio was evaluated using the chi-squared test. To examine the relationship between gut and body length, 59 F3 individuals were used for which genotype segregation was tested, except for one individual for which gut isolation was failed. The significant differences in gut lengths between genotypes were detected using a Bayesian GLM, performed with the brms R package with the default MCMC setting on RStudio ver.1.3. The gut length was modeled with a gamma error structure and log link function. Fixed effects included the standard length and sex in each analysis. For immunohistological analysis, 5 µm serial tissue sections from wild types and mutants (n = 5 each) were prepared using our previously described method (*63*). The sections were incubated with a mouse monoclonal anti-phosphorylated neurofilament H (BioLegend, #smi-31, 1:1000) of which cross-reactivity for medaka had been confirmed in a previous study (*64*). Quantitative analysis of an immunohistological-staining signal was performed using ImageJ software (*36*). We obtained an average of six images of intestinal villus per individual at a 100× objective using light microscopy (BX63, Olympus) equipped with a digital camera (DP74, Olympus). We then split each image into red, green, and blue using the color channels option and subtracted green from the red image with the image calculator. Setting the same threshold, we measured the proportion of the stained area for one intestinal villus of each individual. After standardizing the percentage of the stained area in each experiment, we evaluated the difference of the immunohistological-staining signals between the wild types and the mutants using the Wilcoxon rank-sum test (*P* < 0.05).

### RT-qPCR

We performed reverse transcription-quantitative PCR (RT-qPCR) analyses to evaluate the gene expressions of the *Plxnb3* and *Ppp3r1* genes. We sampled three individuals from each season in the Kabe river (12 individuals totally) for the *Plxnb3* gene expression, and four individuals from four geographical populations (Kamikita and Yokote from NJPN1, Amino from NJPN2, Ichinoseki and Iwata from SJPN1, and Izumi from SJPN4) for the *Plxnb3* and/or the *Ppp3r1* gene expression. The guts were isolated from ethanol-fixed Kabe river samples stored at -20°C and wild-derived laboratory stocks in winter (29 Jan. 2018). The gut was homogenized using a BioMasher (Nippi), and total RNA was isolated from each sample using a combination of the TRIzol (Invitrogen) and NucleoSpin RNA (Macherey-Nagel).

cDNA, which was used as a template for qPCR, was synthesized using a PrimeScript RT reagent kit with gDNA Eraser (TaKaRa Bio) according to the manufacturer’s protocol. *Ribosomal protein L7* (*Rpl7*) was used as an internal reference gene because medaka *Rpl7* was expressed with the less variance among the same tissue samples, in different tissues and stages of development (*65*), and between winter and summer (*66*). The negative control was treated with RNase free water and RT minus products. The reaction mixture consisted of 2 μl of cDNA template, 10 μl of TB Green® Premix Ex Taq™ (Tli RNaseH Plus) (TaKaRa Bio), 0.4 μl of forward and reverse primers (10 µM), and 7.2 μl of RNase-free water. The reaction procedure in a LightCycler® 480 Real-Time PCR System (Roche Diagnostic) was: 95°C, 30 sec; 95°C, 5 sec + 60°C, 30 sec (for amplification, 40 cycles); 95°C, 5 sec + 60°C, 60 sec + 97°C, 0.11°C/sec (for the melting curve); 40°C, 30 sec (for cooling). The primer pairs were: oPlxnb3ex32F1:TGGTGAAGAGCAGTGAAGATCC; oPlxnb3ex33R1:TAGTGTTCCCTTCATGGACAGC for *Plxnb3* (PCR efficiency: 105%), oCANB1ex4F1:TAAACATGAAGGGAAGGCTGGAC; oCANB1ex3R1:ATTCGCCTTCAGGATCTACGAC, for *Ppp3r1* (PCR efficiency: 95%), and oRPL7ex3-4F1:ATCCGAGGTATCAACGGAGTC; oRPL7ex5R1:TGCCGTAGCCACGTTTGTAG for *Rpl7* (PCR efficiency: 95%). The specificity of all PCR primer pairs was confirmed as a single peak in the melting curve analysis. Two technical replicates were run for the RT-qPCR analyses, and the relative expressions of each sample were calculated using the ΔΔCt method with the LightCycler® (Roche Diagnostic). The statistical tests were performed using R ver.4.1.1 and RStudio ver.1.4.1717.

### Outdoor breeding experiment in artificial environment

To test whether the food source affects the gut length, we transported the wild medaka from the Kabe river into the outdoor breeding facility on the Kashiwa campus, The University of Tokyo, in August 2015. Then in September 2016 and February 2017, we sampled and compared the gut lengths of bred and wild medakas in the outdoor breeding facility and the Kabe river, respectively. Their gut lengths were measured according to the method described above.

### Common garden experiment

To examine whether NJPN1 medaka show a shorter gut in winter than in summer, we sampled and measured the medaka gut lengths on 3 February and 30 August 2021 using available wild-derived laboratory stocks. The sampled stocks were: NJPN1: Yokote (n = 10, 10), Miyazu (n = 10, 10), Kaga (n = 7, 10); NJPN2: Kumihama (n = 9, 10), Amino (n = 10, 10), Toyooka (n = 11, 8); SJPNs as control: Ichonoseki (n = 8, 10), Okayama (n = 10, 5), Mishima (n = 10, 9). The significant differences between seasons were detected by the Steel-Dwass test implemented in an NSM3 R package. We used the Bayesian GLMM to detect the gut length variations using the brms R package, as described above. Fixed effects included the standard length, sex, season (winter and summer), genetic background, and the interaction between season and genetic background. Random effects were subject identity and the sampling year. Four MCMC chains were run for 2,000 samples, with a 1,000 sample burn-in, by using every sample. This gave a total of 4,000 samples used. R-hat is the Germany-Rubin convergence statistic for estimating the degree of convergence of a random Markov chain and indicates insufficient convergence at values greater than 1.1.

### Principal component analysis

To test whether or not the population structure had changed in the Kabe river between winter and summer, we performed a principal component analysis (PCA) using the SNPRelate program (*67*) in R ver.3.2.2. A total of 2,167,650 bi-allelic SNPs were generated from the bam files used to detect the methylation regions by MBD-seq. The procedure was such that the SNPs were extracted using SAMtools v1.9 (*40*) mpileup and filtered vcfutils.pl with varFilter -d 6 option, and then only bi-allelic sites were extracted using VCFtools (*57*). The PCA graph was drawn using ggplot2 (*68*).

### Calculation of *d*_A_ distances between seasons in each river based on mtDNA D-loop sequences

The genetic distance matrix was obtained by the calculation made from the number of base differences per site and from the estimation of the net average between groups of mtDNA D-loop sequences using the MEGA7 (*69*). The D-loop sequences were generated by the method previously described (*70*).

These nucleotide sequences were deposited into the international DNA database DDBJ/EMBL/GenBank (accession nos. LC719344 – LC719462).

### Relationships between the number of CpG sites on the upstream of *Plxnb3* and the mutation in the *Ppp3r1* region

To examine the genetic variation on the upstream of *Plxnb3* among geographic populations, we performed PCR direct sequencing as described previously (*35*). The two primers were o*Plxnb3*CpG1F1:GTTCATTTAAGGGAGGAACCAAAGG and o*Plxnb3*CpG1R1:CTCCACTGTGTCATCAAAAGAAGC. Additionally, we determined the *Plxnb3* upstream sequences of related species, *O. curvinotus* and *O. luzonensis* using the shotgun DNA strategy. The libraries were obtained from genomic DNAs extracted in Katsumura et al. 2014 (*35*) using NEBNext URTLA DNA Library Prep Kit (New England Biolabs) and outsourced to Macrogen Japan for sequencing by Hiseq X (Illumina). The short-read data (151 bp paired-end) were filtered using fastp (*50*) with the default setting. The *Plxnb3* upstream sequence extracted from the Japanese medaka Hd-rR genome (ASM223467v1) was used to align the reads using BWA mem (*39*). After the process, the low-quality aligned reads were removed by SAMtools (*40*) and samclip (https://github.com/tseemann/samclip). The *Plxnb3* upstream sequences were obtained using bcftools (*71*) and freebayes (*72*). Finally, The called sites showing nucleotide polymorphisms and insertions/deletions (InDels) were re-examined and conformed by human eyes using the Integrative Genomics Viewer (IGV) (*73*). The short-read data have been submitted to the DRA database under accession number: DRA017652.

*Plxnb3* upstream sequences were aligned across 40 populations and 2 related species, then the CpG sites on these sequences were counted using MEGA X software (*74*). Note that the sequences obtained from the Kaga, Shingu, and Shanghai local populations were excluded from the analysis because their CpG sites of upstream sequences or *Ppp3r1* mutations existed as polymorphisms. Thus, the regression line between the number of CpG sites and gut lengths was calculated using the populations with homozygotes of the C allele on *Ppp3r1*. A phylogenetic tree was reconstructed to estimate the number of CpG sites on ancestral upstream sequences of *Plxnb3* using the maximum likelihood method under the GTR model implemented in MEGA X (*74*). Those numbers were plotted on the medaka genetic group tree obtained from the previous study (*14*), and the allele frequencies of the *Plxnb3* upstream sequence and *Ppp3r1* mutations in each group were calculated as the number of alleles divided by the number of chromosomes. Note that data obtained from the Yongcheon and Shanghai populations were excluded from these analyses because those populations might have experienced artificial gene-flow via the strain maintenance process mentioned in our prior study (*14*).

To test whether the dispersion of the gut length was large, according to the dispersion of the air temperature of each region, we calculated the coefficient of variation (CV) of the gut length and standard deviation of the average annual temperature for each group using the Global Solar Atlas database (https://globalsolaratlas.info). We plotted the standard deviation of the average annual temperature on the X-axis and the mean CV of the gut length on the Y-axis and performed a Pearson’s product-moment correlation test in R and RStudio.

## ACKNOWLEDGMENTS

We thank Mrs. Shizuko Takada, Mrs. Shizuko Chiba, Mrs. Sumiko Tomizuka, Mr. Toshikazu Yamashita, Mrs. Yoshie Sato, Mrs. Ikumi Matsumoto, Mrs. Mayuko Takagi, Mrs. Azusa Matsuda, Mrs. Shiho Kajiwara, Dr. Atsuko Shimada, Prof. Emeritus Akihiro Shima, Prof. Emeritus Nobuo Egami (The University of Tokyo) and Prof. Emeritus Mitsuru Sakaizumi (Niigata University) for maintaining the medaka stocks from wild populations. We also thank NBRP Medaka (https://shigen.nig.ac.jp/medaka/) for providing d-rR/TOKYO (Strain ID: MT837) and accepting our medaka stocks. We are grateful to Mr. Robert E. Brandt, Founder, CEO, and CME, of MedEd Japan, for editing and formatting the manuscript.

## Author contributions

Conceptualization: TK, HO

Methodology: TK, HTakes, YH, YS, HTakeu, HO

Investigation: TK, SS, KY, SO, TG, ST, KF, TN, TI, YY, HTakes, YH, YS

Visualization: TK

Funding acquisition: TK, HO

Project administration: TK

Supervision: SO, HM, MO, HTakeu, HO

Writing – original draft: TK

Writing – review & editing: TK, SO, HTakeu, HO

## Data and materials availability

All data are available in the main text, the supplementary materials or the Dryad repository (*75*). All wild-derived laboratory stocks are available from NBRP medaka (https://shigen.nig.ac.jp/medaka/). The Illumina short-read data and mitochondrial DNA D-loop sequences were deposited to the DNA Data Bank of Japan (DDBJ) (Accession numbers: DRA010581, DRA015161, DRA015162, DRA010605, DRA017652 and LC719344 – LC719462).

## SUPPLEMENTARY MATERIALS

Figs. S1 to S20

Tables S1 to S13

References (*1–76*)

