## Supplementary information for "DNA methylation site loss for plasticity-led novel trait genetic fixation"

**This PDF file includes:**

Figs. S1 to S20
Tables S1 to S13
References

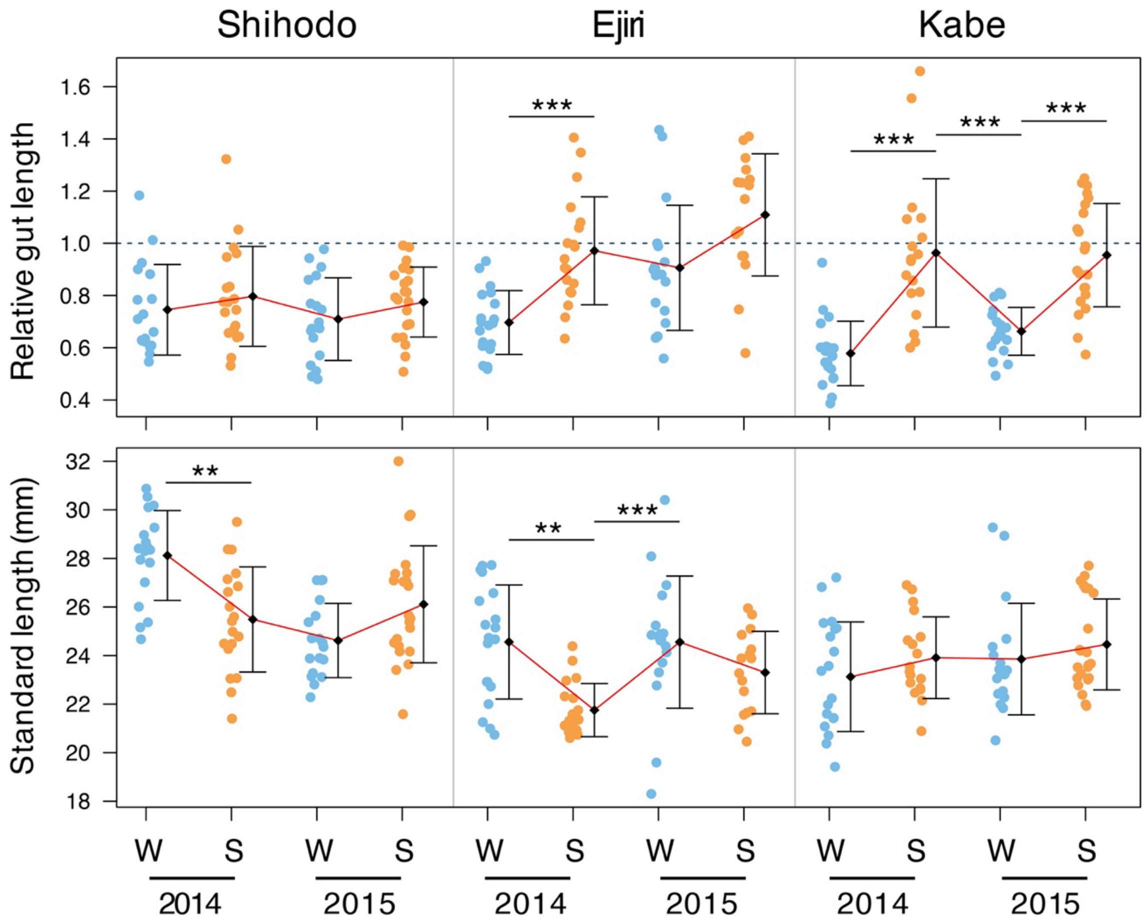


**Fig. S1.** Complete plots of gut length relative to standard length and standard length in medaka from the Shihodo, Ejiri, and Kabe rivers. Relative gut length was calculated by dividing gut length with the standard length. The black diamonds and error bars are the mean of those values and the standard deviation, respectively. The red line joins the mean in each season representing the seasonal variation. A significant difference in relative gut and standard length between seasons was detected using the Steel-Dwass test implemented in the NSM3 R package (***P* < 0.01, ****P* < 0.001)

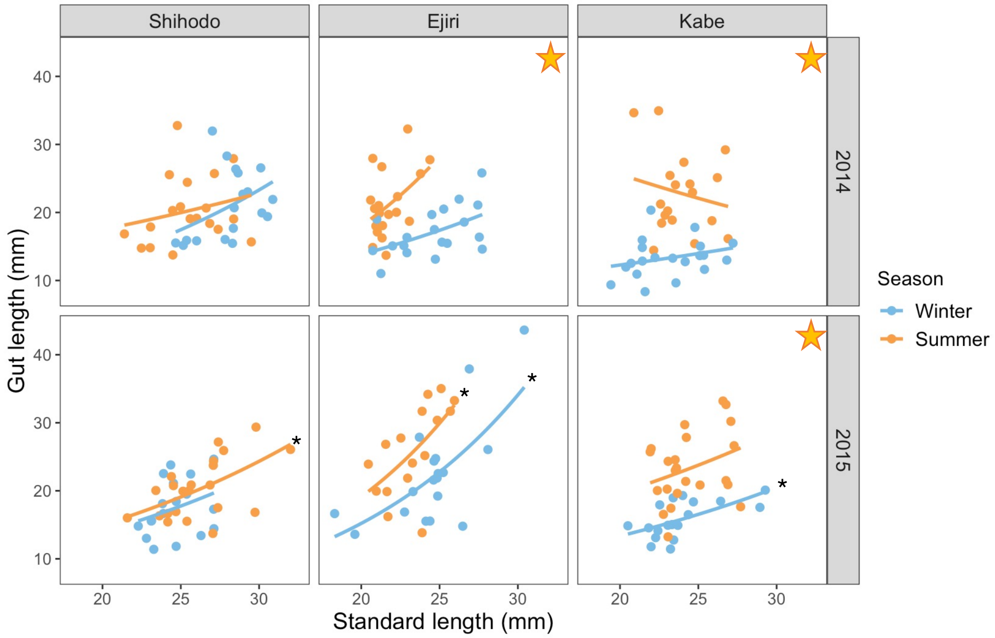


**Fig. S2.** Complete plots of gut length and standard length in each population in Kagawa. A star in a panel indicates that relative gut length differed significantly between winter and summer for that particular river and year (see also Fig. S1). Lines are fitted values from the generalized linear regression with a Gamma distribution. Asterisks indicate the significant correlations between gut and standard length found in that season. The significant correlation coefficients (Spearman's *rho*) were: Shihodo in the summer, *rho* = 0.579; Ejiri in the winter and summer, *rho* = 0.565 and 0.693, respectively; Kabe in the winter, *rho* = 0.666; all in 2015.

**A**

$d_A$ distance (lower left) and standard error (upper right)

| | Ejiri (14 W) | Ejiri (14 S) | Kabe (14 W) | Kabe (14 S) | Shihodo (14 W) | Shihodo (14 S) |
|---|---|---|---|---|---|---|
| Ejiri (14 W) | - | 0.03 | 2.82 | 2.70 | 0.71 | 0.71 |
| Ejiri (14 S) | -0.05 | - | 2.64 | 2.53 | 0.88 | 0.88 |
| Kabe (14 W) | 6.21 | 5.79 | - | 0.06 | 2.96 | 2.96 |
| Kabe (14 S) | 5.64 | 5.26 | -0.01 | - | 2.85 | 2.85 |
| Shihodo (14 W) | 0.99 | 1.05 | 5.61 | 5.04 | - | 0.00 |
| Shihodo (14 S) | 0.99 | 1.05 | 5.61 | 5.04 | 0.00 | - |

**B**

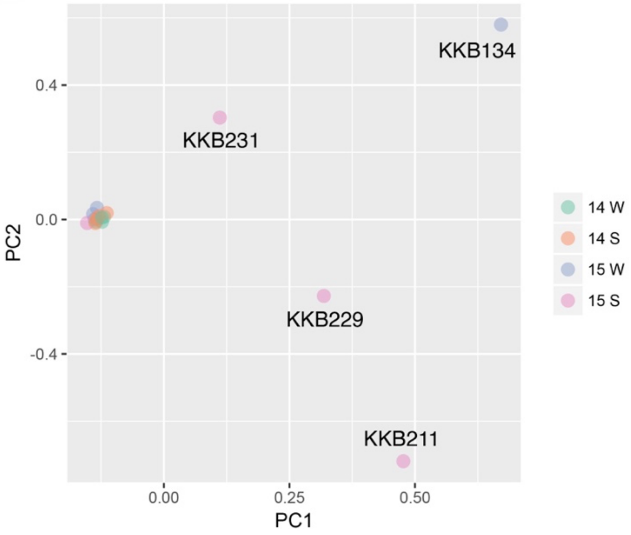


**Fig. S3.** The $d_A$ distance matrix and principal component analysis (PCA). **A**: The genetic distance matrix was based on the number of base differences per site from the estimation of net averages between groups of mtDNA D-loop sequences, generated by the method previously described (*75*). The $d_A$ distances (lower left) and their standard error (upper right) calculated by MEGA7 (*67*) with 1,000 bootstrapping are shown by a factor of 1,000 in the left matrix. The yellow areas in the matrix show the distances and the standard errors obtained between seasons for each population. The matrix indicates only slight population differentiation between seasons in the Ejiri river, suggesting that there was no migration of genetically distant populations among the seasons in the other rivers. **B**: The PCA plot was calculated based on 217 k SNPs generated using the MDB-seq in medaka from the Kabe river. This plot also indicates the no-gene flow among seasons.

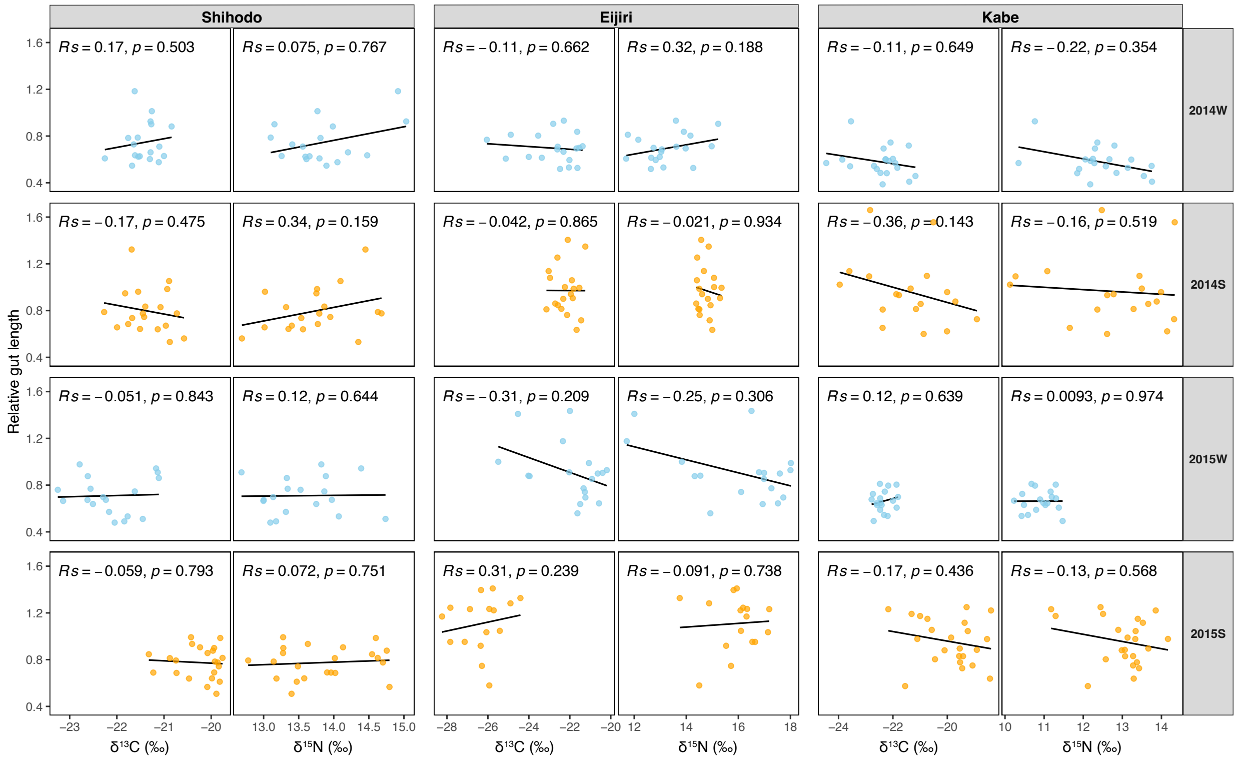


**Fig. S4.** Scatter plots showing the relationship between the stable isotope ratio ($\delta^{13}C$ and $\delta^{15}N$) and relative gut length across three rivers (Shihodo, Ejiri, Kabe) for seasons (2014 Winter/Summer, 2015 Winter/Summer). Stable isotope ratios, $\delta^{13}C$ and $\delta^{15}N$, are indices of a food source diversity and food type (trophic level), respectively. Data points are colored by season: Winter (W) in sky blue and Summer (S) in orange. The solid black line in each panel represents the linear regression fit (y ~ x). The correlation between the variables is assessed using Spearman's rank correlation coefficient (*Rs*) and the corresponding *P*-value, which are displayed within the plot area.

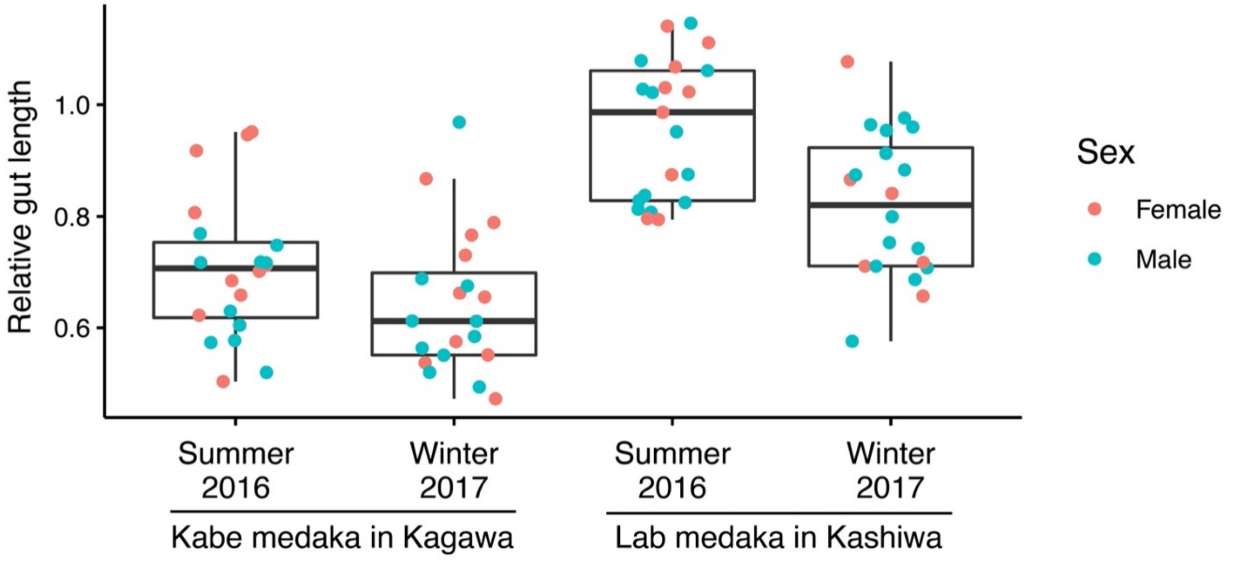


**Fig. S5.** Transplantation experiment of Kabe medaka to the outdoor breeding facility on the Kashiwa campus, The University of Tokyo. We tested whether or not gut plasticity could be observed in medaka collected in the Kabe river kept in a laboratory environment (outdoor breeding facility) and medaka captured in the Kabe river at the same time of year. We confirmed the Kabe medaka showed the seasonal variation of gut length in different environments in Kagawa and Kashiwa using Bayesian GLMM analysis (Table S3), indicating that there is no relationship between the gut length and the food quality.

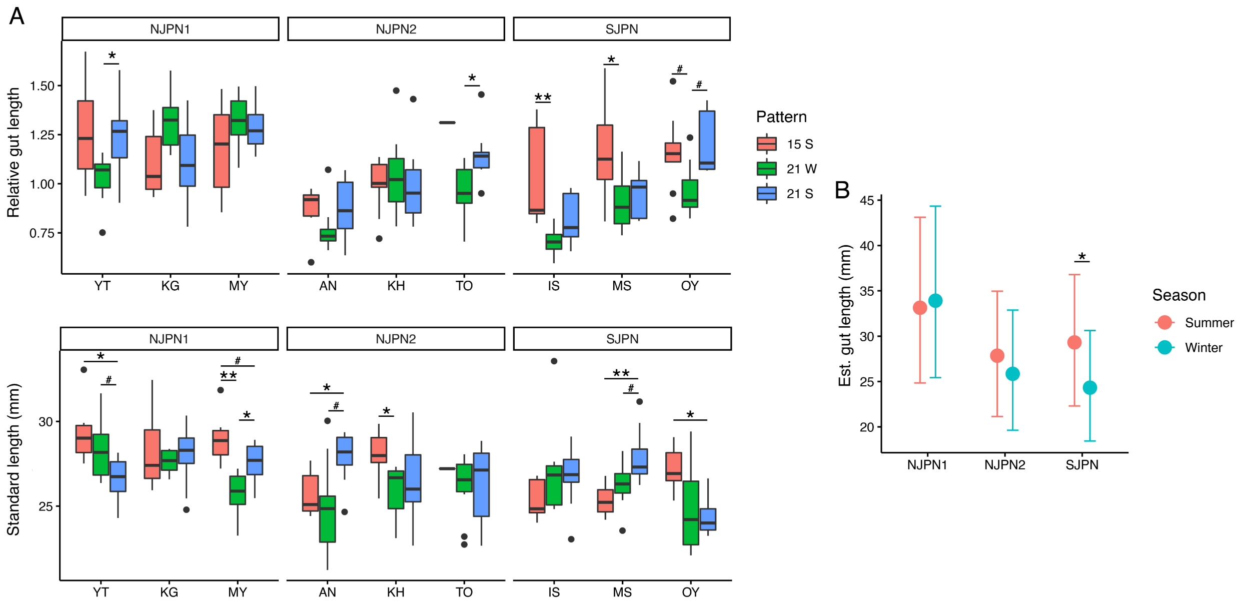


**Fig. S6. A:** Boxplots of gut length relative to standard length and standard length in medaka from NJPN1 (YT, Yokote; KG, Kaga; MY, Miyazu), NJPN2 (AN, Amino; KH, Kumihama; TO, Toyooka) and SJPN (IS, Ichinoseki; MS, Mishima; OY, Okayama). Relative gut length was calculated by dividing gut length with standard length. The colors of boxplots represent year and season (15S: summer in 2015; 21W: winter in 2021; 21S: summer in 2021). The asterisks indicate a significant difference in relative gut and standard length between seasons using the Steel-Dwass test ($^{\#}P < 0.10$, $^{*}P < 0.05$, $^{**}P < 0.01$). **B:** Seasonal gut length changes corrected by standard length in the subgroups. The blue and orange dots, their error bars indicate the mean gut length and 95% CI (credible interval) after adjusting for the effect of body length in winter and summer, respectively, as estimated by Bayesian GLMM. The asterisk indicates that medakas in the winter had significantly shorter guts than did those in the summer (see also Table S6).

| Number of DMRs detected for all possible patterns | | | | | |
|---|---|---|---|---|---|
| DNA methylation pattern | | | | DMRs | |
| 14W | 14S | 15W | 15S | "frequent" DMRs | %FDR |
| | | | | 192 | 16.6 |
| | | | | 14 | >100 |
| | | | | 7567 | 2.8 |
| | | | | 334 | 9.7 |
| | | | | 710 | 15.8 |
| | | | | 71 | >100 |
| | | | | 18 | >100 |
| | | | | 23 | >100 |
| | | | | 69 | >100 |
| | | | | 58 | >100 |
| | | | | 78 | >100 |
| | | | | 73 | >100 |
| | | | | 67 | >100 |
| | | | | 6 | >100 |

**Fig. S7.** Differential methylation regions (DMR) found using MethylAction (*38*). This shows the number of DMRs detected in all combinations of patterns (hyper- [black and gray squares] and hypomethylation [white squares]). The pattern with black squares represents seasonal methylation (wSwS and WsWs). The color code is the same in Figure 2. The "frequent" DMRs correspond to those whose methylation status is agreed in more than two-thirds of the samples within each group (>2/3).

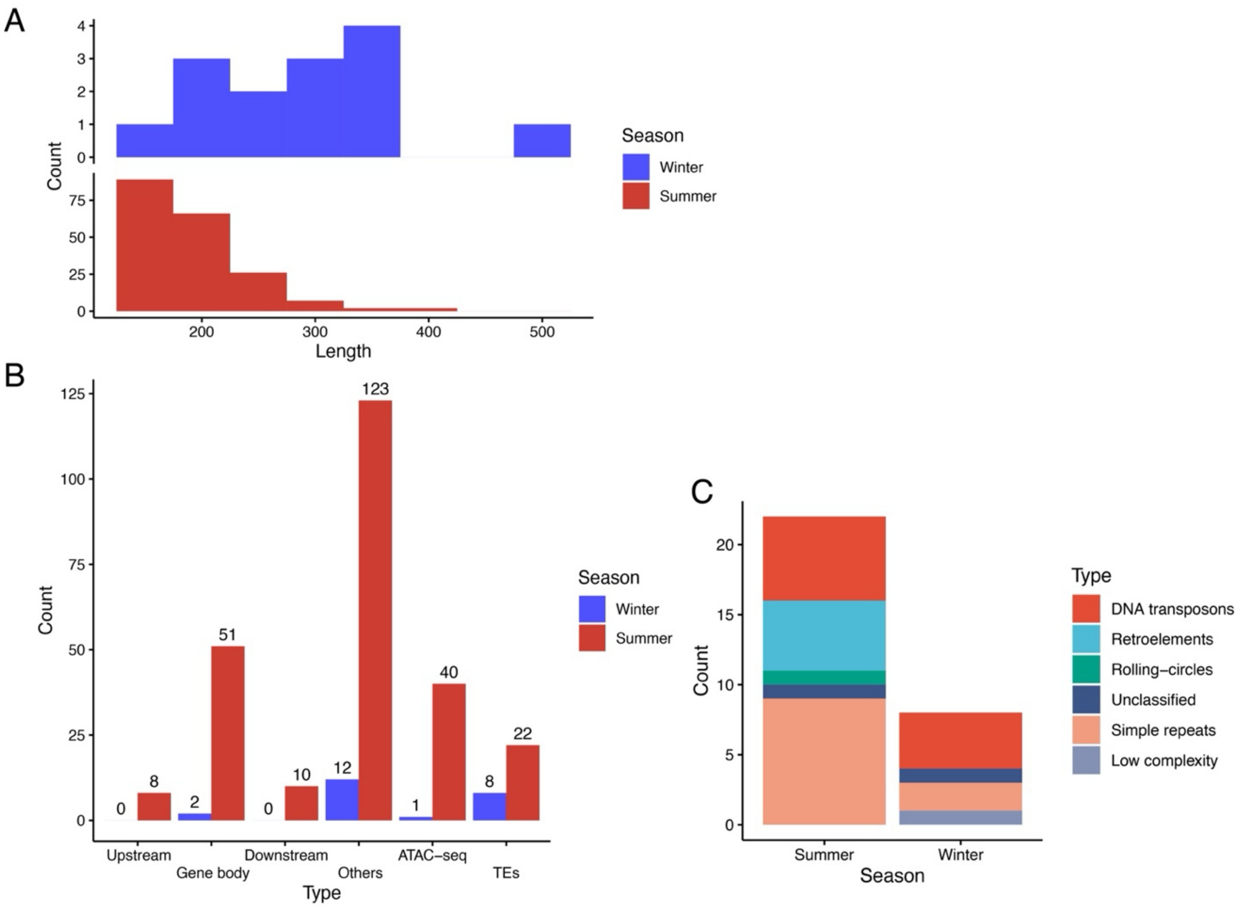


**Fig. S8.** The annotated MBD-seq peak regions. **A**: The length distribution of the regions methylated in summer or winter. **B**: Association of putative seasonally methylated regions with annotated genomic regions. "Upstream" and "Downstream" indicate the regions within 2 kbp from adjusted genes. "Others" indicate distant regions from genes (>2 kbp). **C**: TE-related regions in putative seasonally methylated regions.

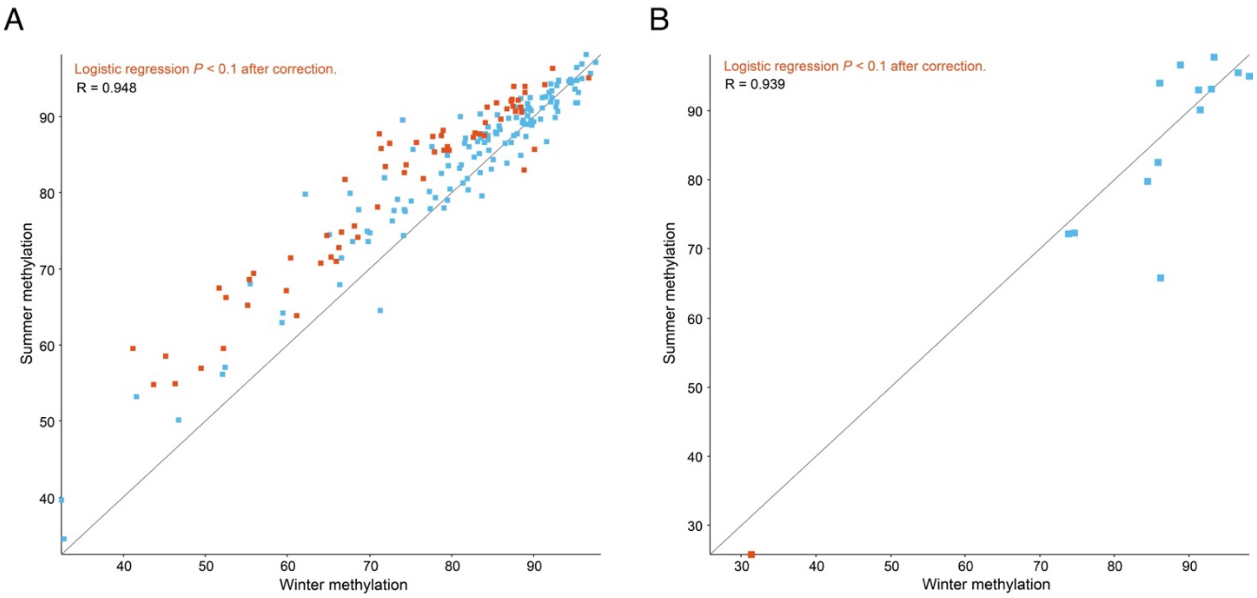


**Fig. S9.** Scatter plots between winter and summer methylation, calculated by WGBS data, on the regions detected by MDB-seq. WGBS-based methylation levels in the summer methylation region (**A**) and the winter methylation region (**B**) detected by MBD-seq are shown. Significant differences were detected in 63/192 and 1/14 regions, respectively. Gray and blue indicate regions detected by MBD-seq and regions supported by logistics regression analysis from WGBS data as significantly different between seasons, respectively ($P < 0.10$ after multiple correction).

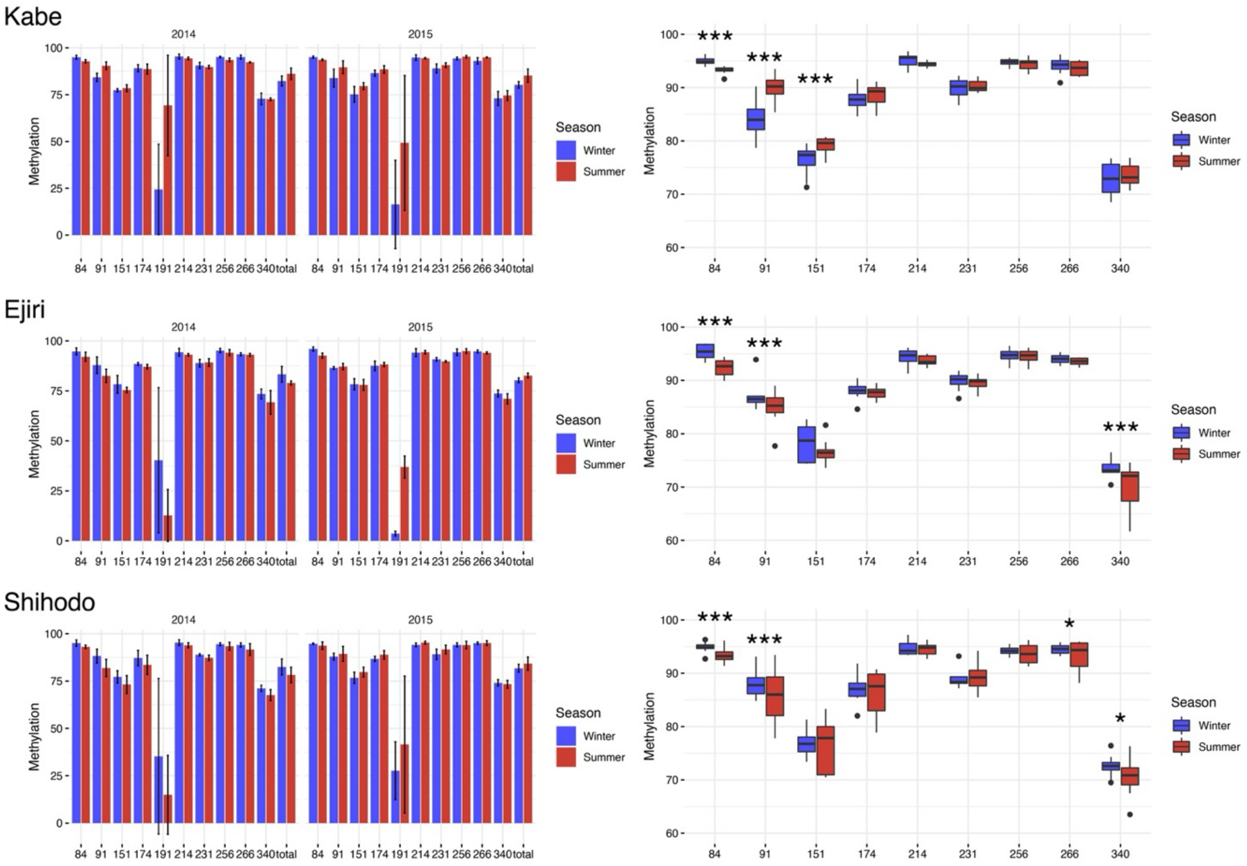


**Fig. S10.** Comparison of methylation between winter and summer on the *plxnb3* upstream region. The barplot shows methylation at each CpG site and the total average including the SNP site (the 5th CpG site presented as the 191st site in the amplified PCR product). The error bar represents the standard deviation. The boxplot shows seasonal methylation (2014 + 2015) at each CpG site exclude the SNP site. The asterisks indicate significant differences in methylation between seasons using logistic regression analysis (*$P < 0.05$, **$P < 0.01$, ***$P < 0.001$).

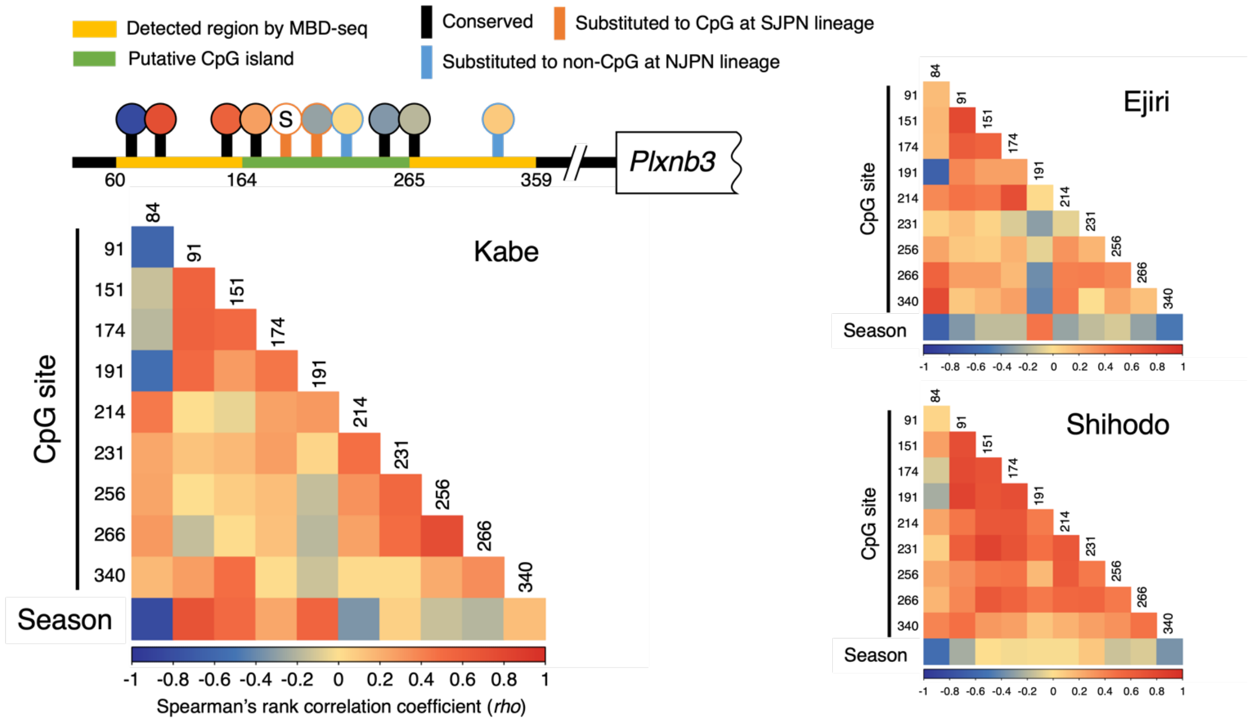


**Fig. S11.** Correlation of methylation between CpG sites and seasons on the *Plxnb3* upstream region in each river population, based on targeted bisulfite sequencing (tBS) data. The upper left figure shows the DMR detected by MBD-seq (yellow), CpG island (yellow-green), the position of the CpG sites (vertical bars), and the position of the SNP site (S). The numbers indicate the nucleotide positions on the amplified region (422 bp) by the tBS. The colors of vertical bars represent the following CpG states: black, conserved among medaka populations; orange, varied in the SJPN lineage; light blue, lost in the NJPN lineage. The color differences in the circles and the heat maps indicate the Spearman's rank correlation coefficient (*rho*) which was calculated between methylations on the CpG sites, and between methylation on each CpG site and season as 0 for winter and 1 for summer. The numbers on the left and top on each row and column of the heatmaps show the nucleotide positions of the CpG sites on the amplified region (422 bp) by the tBS. These heatmaps show that the methylation between neighboring CpG sites is well correlated, and that there are not only seasonally methylated CpG sites but also constantly (stably) methylated CpG sites regardless of seasons in the upstream region of *Plxnb3*.

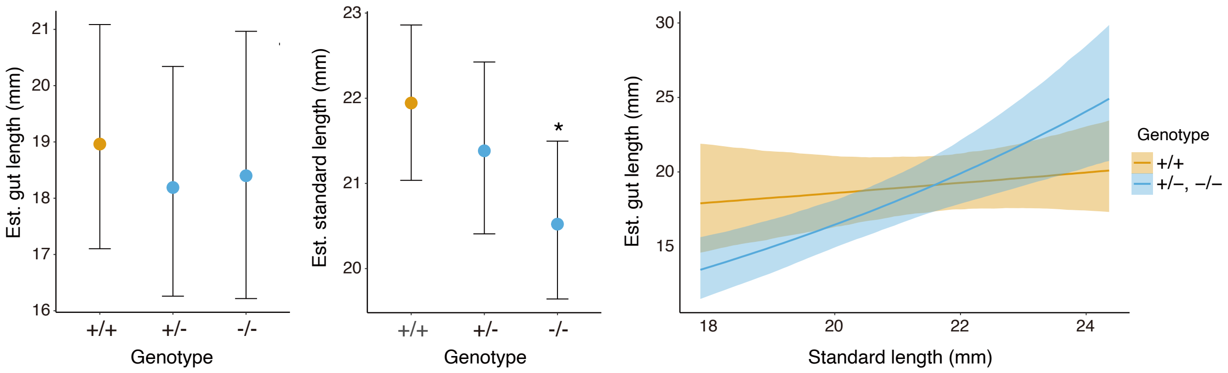


**Fig. S12.** Comparisons of standard and gut lengths among genotypes were performed in the Bayesian generalized linear model (GLM) framework. Points and error bars were fitted values and 95% Bayesian CIs (full data in Fig. 3). The asterisk indicates that -/- exhibited significantly shorter body length than did the other genotypes. The gut length is associated with the standard length in the mutants with deletion, not the wild type. Lines and shaded areas are fitted values and 95% CIs from the Bayesian GLM. The yellow and blue indicate the wild type and mutants (both of the heterozygote and homozygote), respectively. Their estimates by GLM are in table S9.

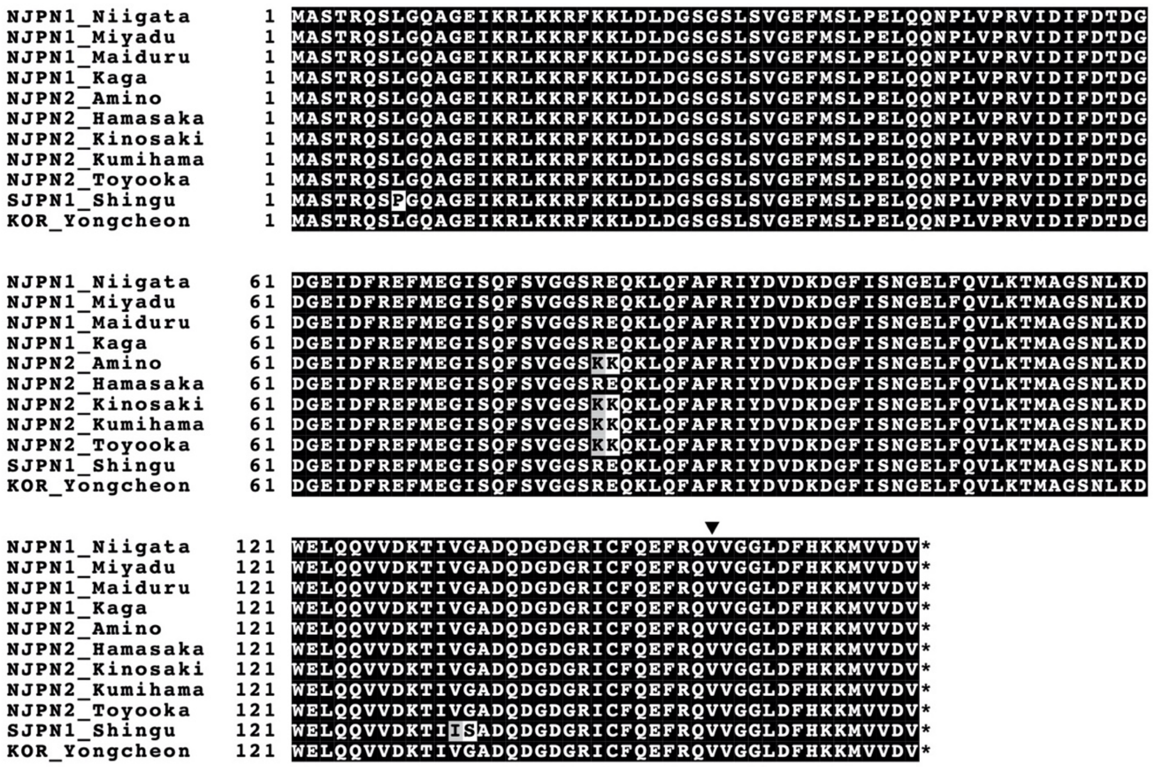


**Fig. S13.** Alignment of the *Ppp3r1* coding sequence region. Eleven sequences were determined using the Sanger method. The black arrowhead shows the position of amino acid encoded by codon in the synonymous SNP detected by RAD-seq. Unique amino acid substitution was not found within the NJPN1 subgroup, indicating the changes in *Ppp3r1* expression (but not in that function) contribute to gut length.

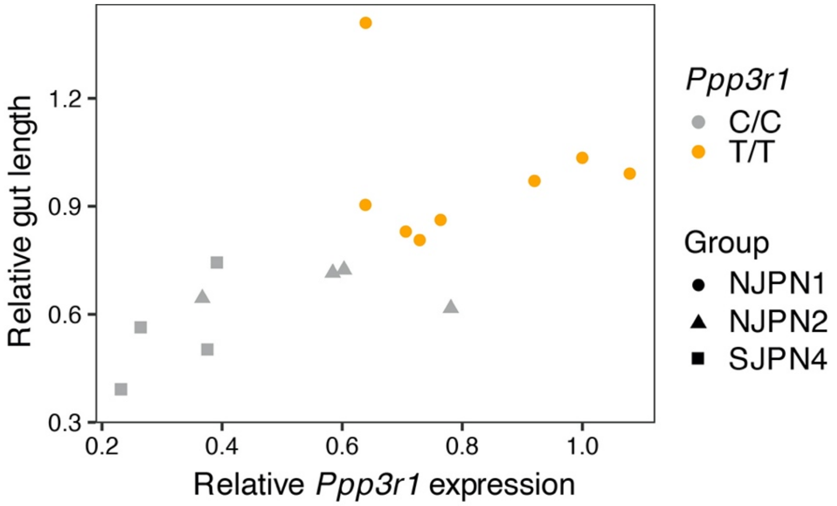


**Fig. S14.** The complete plot of the relative gut length and the relative *Ppp3r1* expression. The GLMM analysis in Fig. 4C was performed on this data irrespective of *Ppp3r1* genotype. We analyzed the data in two ways: (1) without controlling for genetic background, and (2) with genetic background included as a random effect (see also table S11).

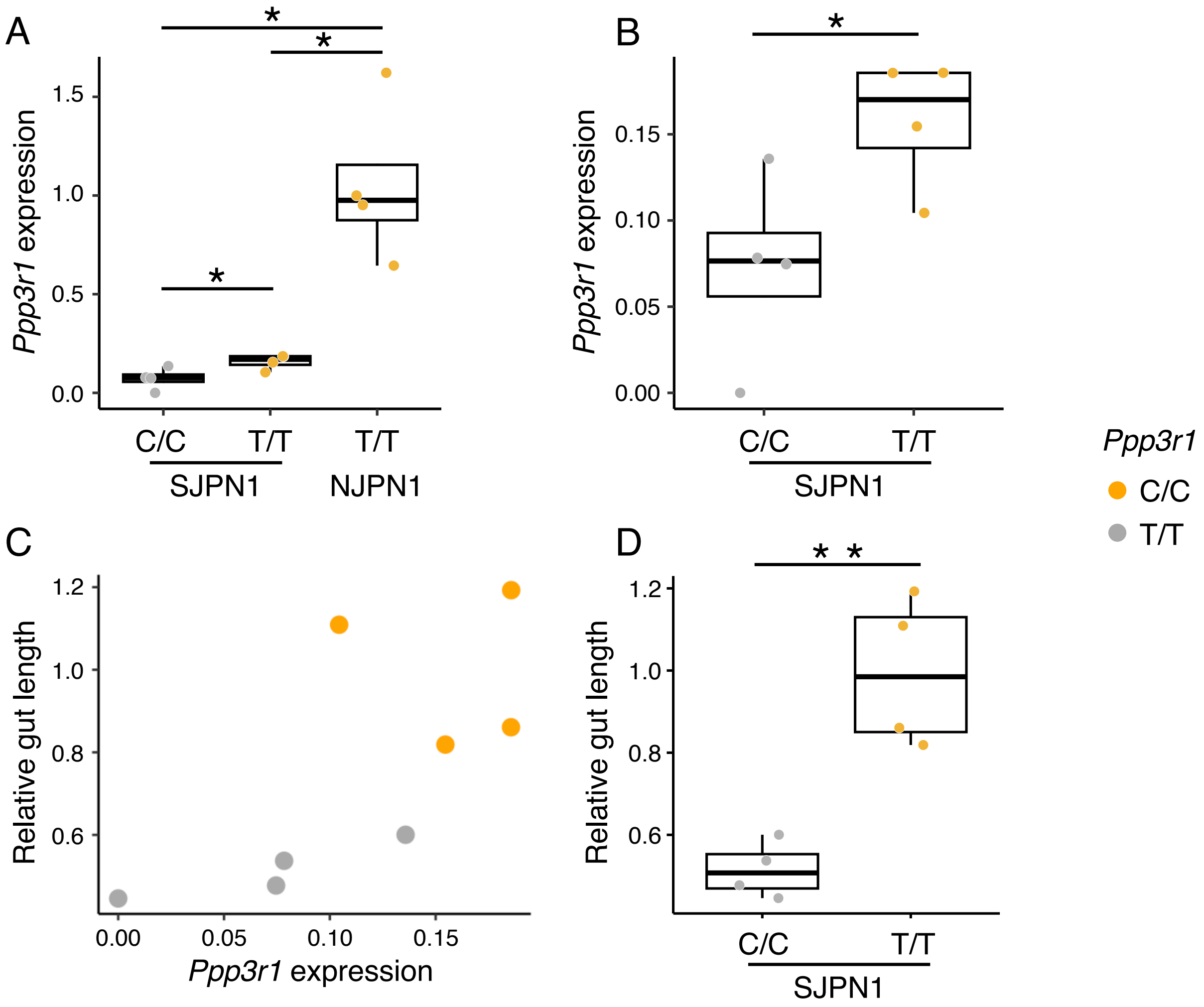


**Fig. S15.** Boxplots of the relative *Ppp3r1* expression among *Ppp3r1* genotypes in the same genetic background, SJPN1, with NJPN1 (**A**) and without NJPN1 (**B**); a scatterplot of the relative gut length and relative *Ppp3r1* expression using SJPN1 medakas (**C**); and a boxplot of the relative gut length between *Ppp3r1* genotypes in SJPN1 **(D)**. The orange and grey points indicate the T/T and C/C genotypes of *Ppp3r1*, respectively. The medakas with these genotypes were obtained from the following populations: the C/C genotype from Ichinoseki, SJPN1; the T/T genotype from Iwata, SJPN1; and the T/T genotype from Yokote, NJPN1. The statistical significance in (**A**) and (**B**) was evaluated using Welch's *t* test with the Holm correction for multiple comparisons (*$P < 0.05$). The correlation in **(C)** was assessed by Spearman's rank correlation coefficient test (*rho* = 0.857, $P = 0.011$). The statistical significance in **(D)** was evaluated using Welch's t-test (**$P < 0.01$).

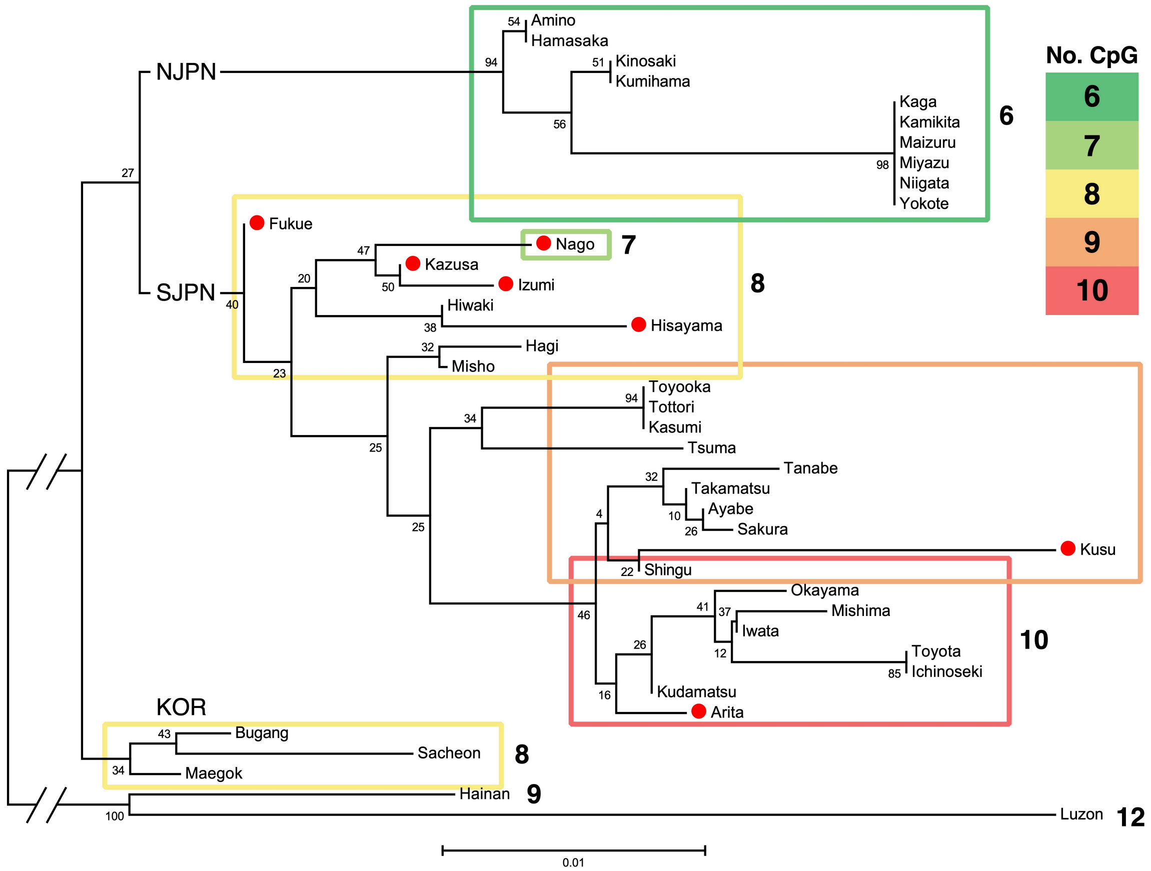


**Fig. S16.** Neighbor-joining tree based on a Jukes-cantor + gamma model (shape parameter = 0.38) using 311 bp sequences of seasonal methylated regions. The frame surrounding the OTUs matches the color key in the upper right corner, indicating the number of CpG sites. The red circles represent northern Kyushu medaka, which is the origin of the SJPN groups. This tree indicates that the variations of the numbers of CpG sites had already existed in northern Kyushu before the spread to the eastern part of Japan, and that CpG sites had rapidly decreased by nucleotide substitutions in the common ancestor of the NJPN groups.

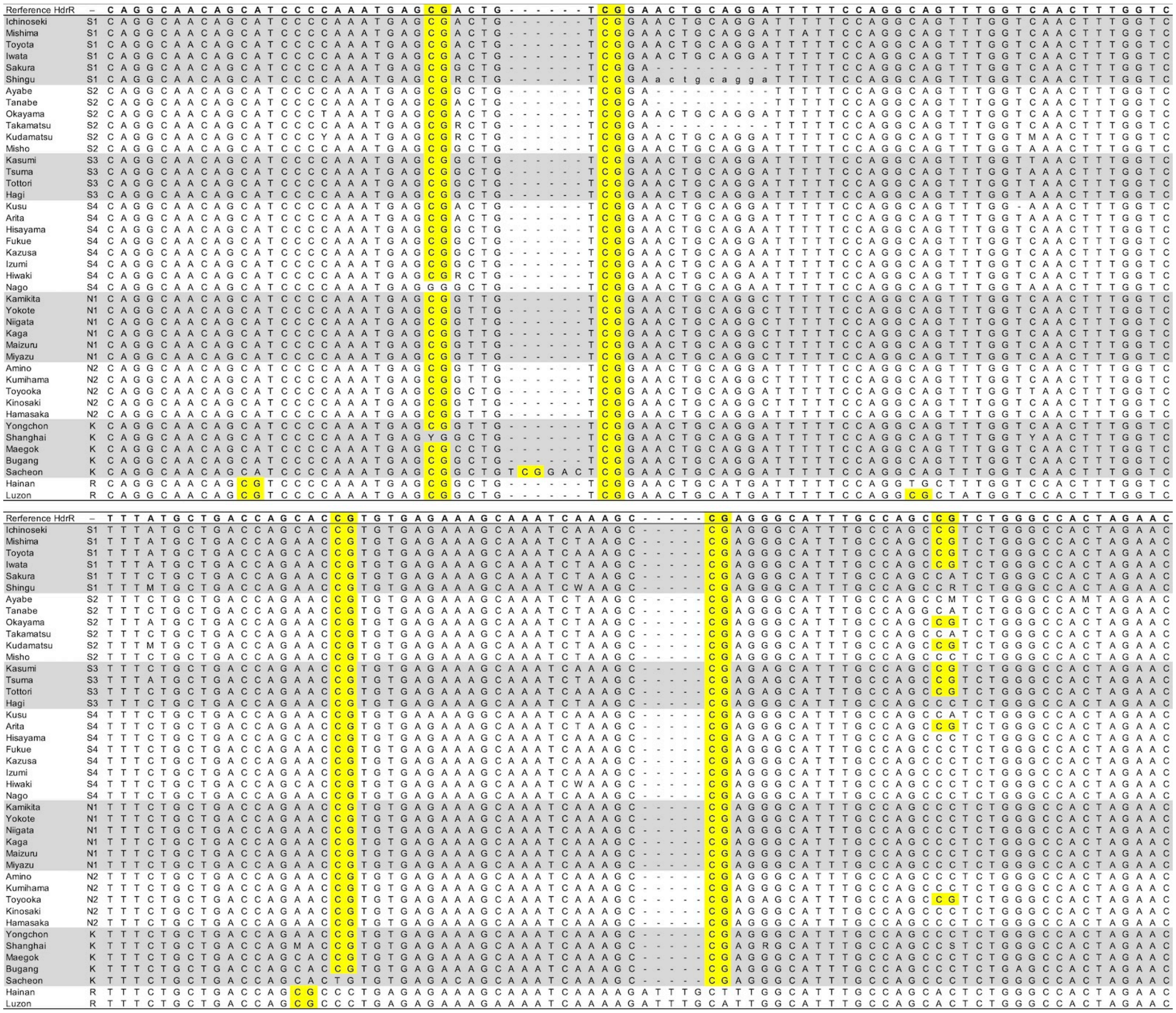

**Fig. S17.** Alignment of *Plxnb3* upstream sequences in the seasonal methylated region shown using MBD-seq analysis. There were 42 sequences determined using the Sanger and the high-throughput sequencing. The top sequence is that extracted from the Hd-rR genome. Each CpG site is highlighted in yellow. The abbreviations (S1, S2, S3, S4, N1, N2, and K) in the second column from the left are the same as those in Figure 1C, and "R" means the related species. The populations are grouped in gray and white backgrounds. The nucleotide sequence marked "---" and small letters indicate deletions and heterozygous indels, respectively.

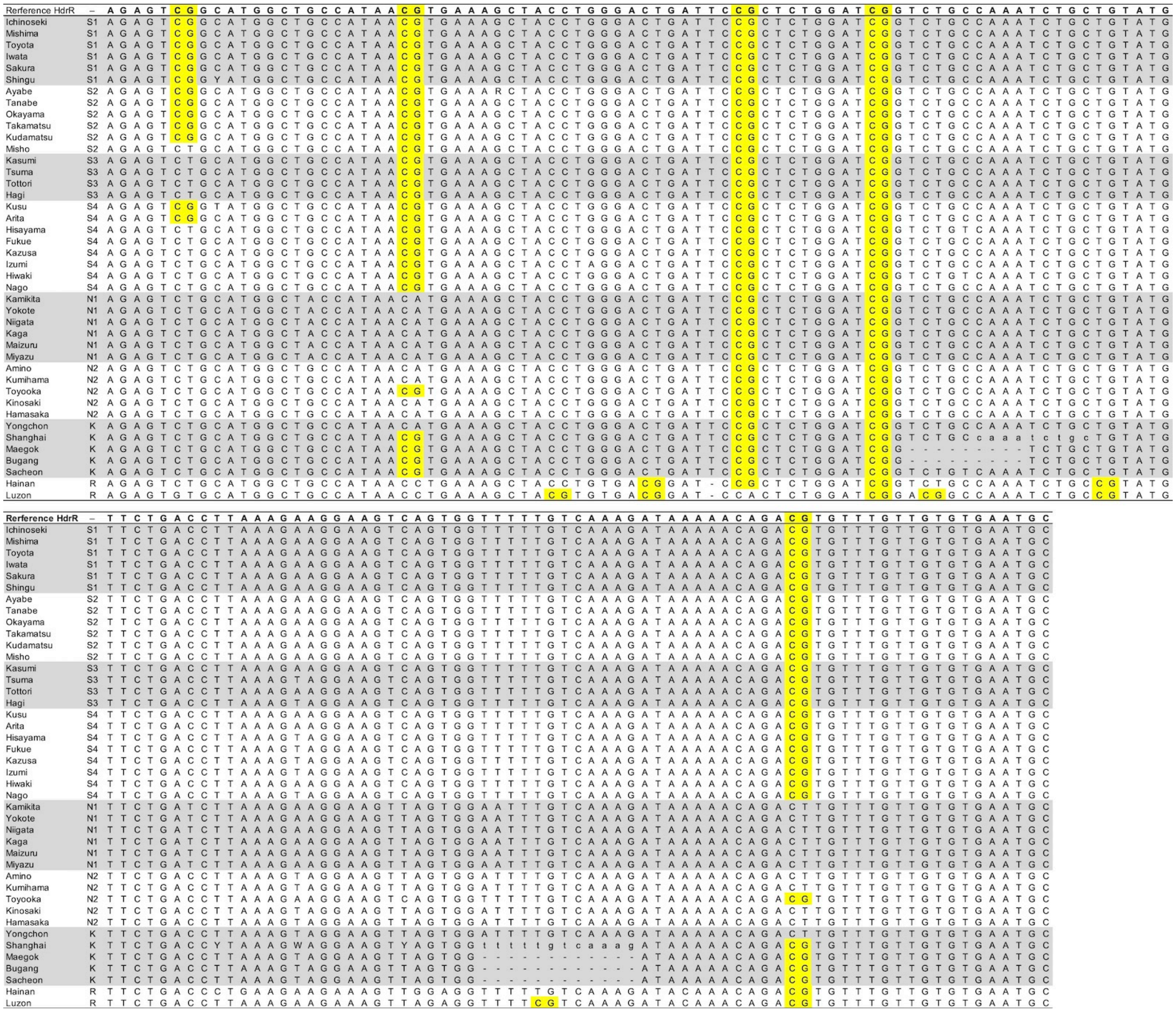

**Fig. S17.** (*continued*)

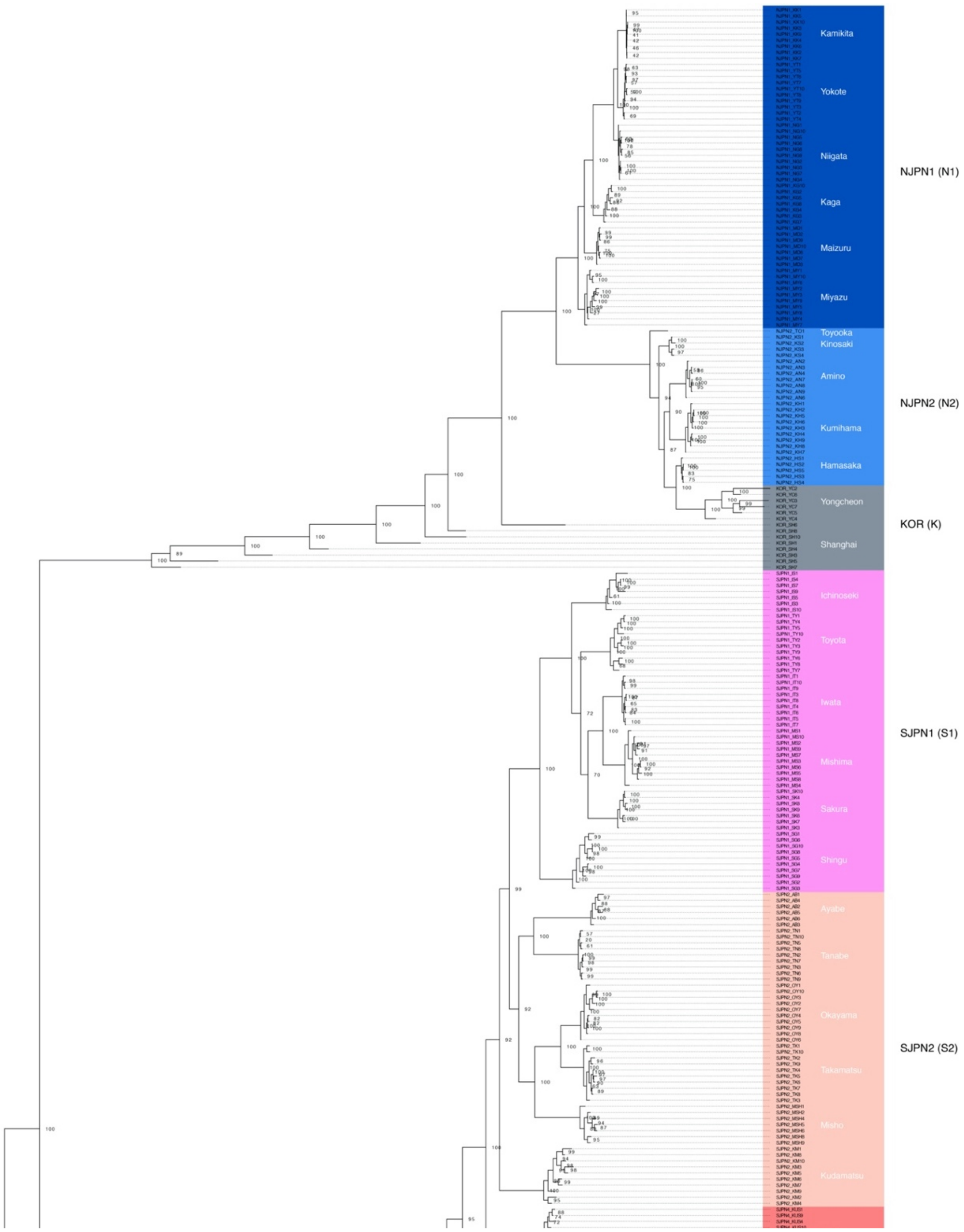


**Fig. S18.** A phylogenetic tree reconstructed by the maximum likelihood method using the general time reversible (GTR) model and the ascertainment bias correction (ASC) with ultrafast bootstrap values as node labels. The colors are the same as those in Fig. 1C, which represent the subgroups in the medaka populations (SJPN4 included Northern and Southern Kyushu subgroups). This tree topology supports our previous study *(14)*, which mentioned that the Yongcheon and Shanghai populations may have experienced artificial gene flow through strain maintenance.

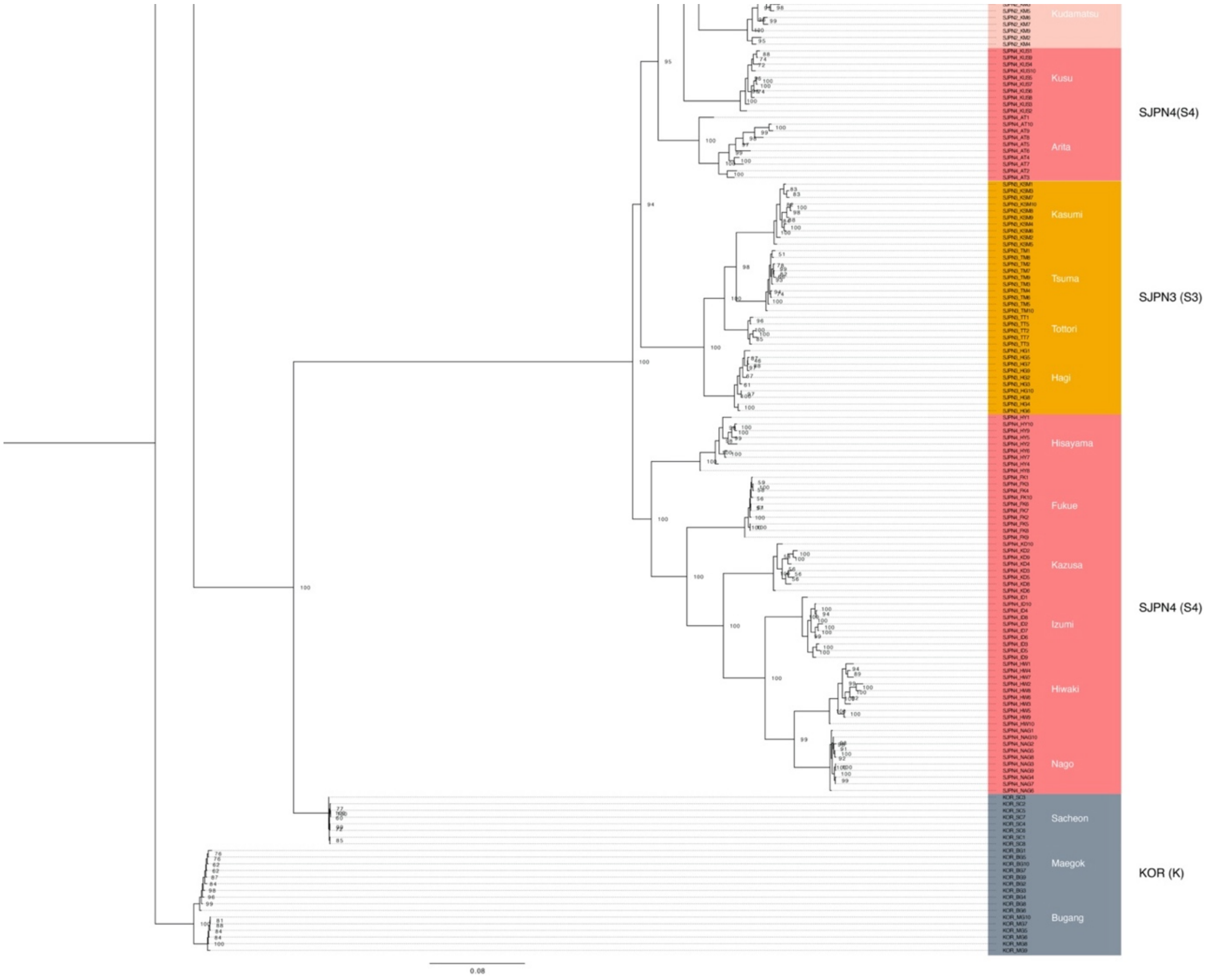


**Fig. S18.** (*continued*)

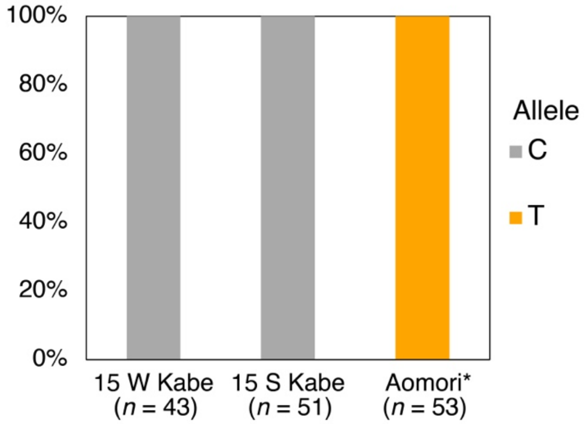


**Fig. S19.** The *Ppp3r1* allele frequency in the wild populations. The gut-length associated T-allele was fixed in the Aomori populations (NJPN1), and the other C-allele was fixed in the winter (15 W Kabe) and the summer (15 S Kabe) in the Kabe populations in 2015. The DNA samples for genotyping the Aomori population were the same as those in Katsumura *et al.* (*76*).

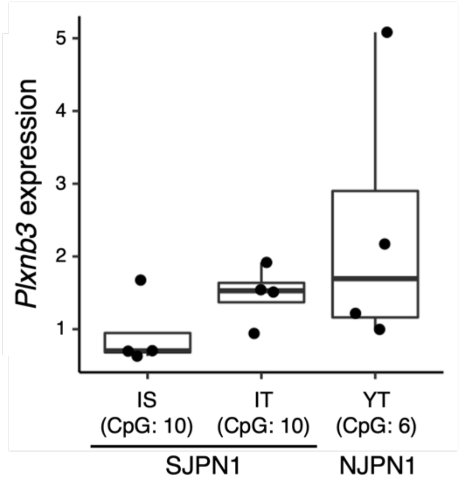


**Fig. S20.** A boxplot of the relative *Plxnb3* expression among the populations. The number in the parentheses shows the number of CpG sites in the seasonal methylated region. The capital letters are the following: IS, Ichinoseki from SJPN1; IT, Iwata from SJPN1; YT, Yokote from NJPN1. The *Plxnb3* expression showed the uneven variance among the populations, which was detected using the Bartlett test ($P < 0.05$).

**Table S1.** The number of individuals sampled from each site in Kagawa.

| | 2014 | | 2015 | | |
|---|---|---|---|---|---|
| | 24–26 Jan. | 29–31 Aug. | 20–22 Feb. | 28–30 Aug. | |
| Rivers | Winter | Summer | Winter | Summer | Total |
| Kabe | 20 | 18 | 18 | 23 | 79 |
| Ejiri | 19 | 19 | 18 | 16 | 72 |
| Shihodo | 18 | 19 | 18 | 22 | 77 |
| Total | 57 | 56 | 54 | 61 | 228 |

**Table S2**. Parameter estimates for fixed and random effects from Bayesian GLMM models of gut lengths in medaka from the Kabe, Ejiri, and Shihodo rivers, with individual ID included as a random effect to account for unmeasured individual variation. Effect sizes of winter and male are given relative to the model intercepts (summer and female, respectively) and are the posterior means of 100,000 sampled iterations. The lower (l) and upper (u) credible intervals (CIs) are given at the 95% significance level. Four MCMC chains were run for 105,000 samples, with a 5,000 sample burn-in, followed by every 4th sample being used. This gave a total of 100,000 samples used. R-hat is the Gelman-Rubin convergence statistic for estimating the degree of convergence of a random Markov chain and indicates insufficient convergences at values greater than 1.1. The effects when the 95% CI did not include zero are indicated in bold.

The Kabe river

| Fixed effects | Estimate | Est. Error | l-95% CI | u-95% CI | R-hat |
|---|---|---|---|---|---|
| **Intercept** | **2.51** | **0.32** | **1.88** | **3.13** | **1.00** |
| Sex: male | -0.06 | 0.05 | -0.16 | 0.04 | 1.00 |
| Standard length | 0.03 | 0.01 | 0 | 0.05 | 1.00 |
| **Season: Winter** | **-0.44** | **0.05** | **-0.54** | **-0.34** | **1.00** |

| Random effect | Estimate | Est. Error | l-95% CI | u-95% CI | R-hat |
|---|---|---|---|---|---|
| **ID** | **0.12** | **0.06** | **0.01** | **0.23** | **1.00** |

The Ejiri river

| Fixed effects | Estimate | Est. Error | l-95% CI | u-95% CI | R-hat |
|---|---|---|---|---|---|
| **Intercept** | **1.5** | **0.3** | **0.91** | **3.13** | **1.00** |
| Sex: male | -0.08 | 0.06 | -0.18 | 0.04 | 1.00 |
| **Standard length** | **0.07** | **0.01** | **0.05** | **0.05** | **1.00** |
| **Season: Winter** | **-0.33** | **0.06** | **-0.45** | **-0.34** | **1.00** |

| Random effect | Estimate | Est. Error | l-95% CI | u-95% CI | R-hat |
|---|---|---|---|---|---|
| **ID** | **0.12** | **0.07** | **0.01** | **0.24** | **1.00** |

The Shihodo river

| Fixed effects | Estimate | Est. Error | l-95% CI | u-95% CI | R-hat |
|---|---|---|---|---|---|
| **Intercept** | **1.94** | **0.31** | **1.34** | **2.54** | **1.00** |
| Sex: male | -0.05 | 0.05 | -0.16 | 0.06 | 1.00 |
| **Standard length** | **0.04** | **0.01** | **0.02** | **0.06** | **1.00** |
| Season: Winter | -0.07 | 0.05 | -0.17 | 0.03 | 1.00 |

| Random effect | Estimate | Est. Error | l-95% CI | u-95% CI | R-hat |
|---|---|---|---|---|---|
| **ID** | **0.13** | **0.06** | **0.01** | **0.22** | **1.00** |

**Table S3.** Parameter estimates for fixed and random effects from Bayesian GLMM models of gut lengths in the transplantation experiment (fig. S5), with individual ID and breeding environment (Kagawa or Kashiwa) included as random effects to account for unmeasured individual and environmental variation, respectively. Effect size of winter and male are given relative to the model intercepts (summer and female, respectively). The lower and upper CIs are given at the 95% significance level. Four MCMC chains were run for 105,000 samples, with a 5,000 sample burn-in, followed by every 4th sample being used. This gave a total of 100,000 samples used.

| Fixed effects | Estimate | Est. Error | l-95% CI | u-95% CI | R-hat |
|---|---|---|---|---|---|
| **Intercept** | **2.19** | **0.77** | **0.55** | **3.81** | **1.00** |
| **Standard length** | **0.03** | **0.01** | **0.01** | **0.06** | **1.00** |
| Sex: male | -0.07 | 0.04 | -0.16 | 0.01 | 1.00 |
| **Season: winter** | **-0.16** | **0.06** | **-0.27** | **-0.05** | **1.00** |

| Random effects | Estimate | Est. Error | l-95% CI | u-95% CI | R-hat |
|---|---|---|---|---|---|
| **ID** | **0.09** | **0.05** | **0.01** | **0.17** | **1.00** |
| **Environment** | **0.96** | **1.01** | **0.11** | **3.74** | **1.00** |

**Table S4:** Statistics of wild lab-stocks used in this study. The asterisks indicate northern Kyushu populations.

| Group | Subgroup | Population (NBRP ID) | Location number in Katsumura *et al.* (*14*) | Sampling date | N | Average length (mm) | |
|---|---|---|---|---|---|---|---|
| | | | | | | Body | Gut |
| NJPN | NJPN1 | Kamikita (WS1590) | 1 | 28Sep2015 | 9 | 28.69 | 32.94 |
| | | Yokote (WS1593) | 4 | 24Sep2015 | 10 | 29.19 | 37.15 |
| | | Niigata (WS1596) | 7 | 26Sep2015 | 10 | 28.51 | 38.29 |
| | | Kaga (WS1599) | 10 | 21Sep2015 | 7 | 28.3 | 31.48 |
| | | Maizuru (WS1602) | 13 | 20Oct2015 | 10 | 25.86 | 29.82 |
| | | Miyazu (WS1603) | 14 | 27Aug2015 | 10 | 28.92 | 33.95 |
| | NJPN2 | Amino (WS1660) | 21 | 2Sep2015 | 7 | 25.75 | 22.3 |
| | | Kumihama (WS1661) | 22 | 31Aug2015 | 9 | 27.9 | 27.54 |
| | | Toyooka (WS1662) | 23 | 2Sep2015 | 1 | 27.21 | 35.68 |
| | | Kinosaki (WS1663) | 24 | 31Aug2015 | 4 | 27.17 | 24.81 |
| | | Hamasaka (WS1664) | 25 | 2Sep2015 | 5 | 26.48 | 21.31 |
| SJPN | SJPN1 | Ichinoseki (WS1605) | 27 | 6Oct2015 | 7 | 25.43 | 26.61 |
| | | Toyota (WS1612) | 34 | 7Oct2015 | 10 | 29.24 | 29.3 |
| | | Mishima (WS1618) | 40 | 5Oct2015 | 10 | 25.39 | 29.53 |
| | | Iwata (WS1620) | 42 | 9Oct2015 | 9 | 29.64 | 41.88 |
| | | Sakura (WS1609) | 31 | 7Oct2015 | 7 | 29.41 | 26.57 |
| | | Shingu (WS1623) | 45 | 21Oct2015 | 10 | 25.79 | 23.9 |
| | SJPN2 | Ayabe (WS1629) | 61 | 20Oct2015 | 6 | 30.87 | 30.67 |
| | | Tanabe (WS1642) | 64 | 25Sep2015 | 9 | 27.64 | 24.2 |
| | | Okayama (WS1644) | 66 | 16Oct2015 | 10 | 27.13 | 31.42 |
| | | Takamatsu (WS1647) | 69 | 16Oct2015 | 10 | 28.3 | 32.58 |
| | | Kudamatsu (WS1651) | 73 | 15Oct2015 | 10 | 25.51 | 31.74 |
| | | Misho (WS1625) | 47 | 14Oct2015 | 7 | 25.59 | 22.85 |
| | SJPN3 | Kasumi (WS1654) | 76 | 18Sep2015 | 10 | 27.88 | 29.74 |
| | | Tsuma (WS1658) | 80 | 15Sep2015 | 10 | 31.38 | 24.48 |
| | | Tottori (WS1656) | 78 | 15Sep2015 | 5 | 28.46 | 34.82 |
| | | Hagi (WS1659) | 81 | 14Sep2015 | 10 | 27.65 | 24.88 |
| | SJPN4 | Kusu* (WS1627) | 49 | 19Oct2015 | 10 | 25.57 | 23.13 |

| | | | | | | | |
|---|---|---|---|---|---|---|---|
| | | Arita* (WS1632) | 54 | 22Oct2015 | 10 | 28.13 | 23.06 |
| | | Hisayama* (WS1631) | 53 | 14Sep2015 | 9 | 29.38 | 26.92 |
| | | Fukue* (WS1630) | 52 | 28Sep2015 | 10 | 28.65 | 30.81 |
| | | Kazusa* (WS1633) | 55 | 29Sep2015 | 9 | 30.02 | 35.03 |
| | | Izumi (WS1634) | 56 | 2Oct2015 | 10 | 28.89 | 22.3 |
| | | Hiwaki (WS1635) | 57 | 2Oct2015 | 10 | 26.84 | 25.95 |
| | | Nago (WS1637) | 59 | 5Oct2015 | 10 | 28.55 | 27.83 |
| KOR/CHN | KOR | Yongcheon (WS1668) | 18 | 25Aug2015 | 6 | 29.79 | 21.77 |
| | | Maegok (WS1666) | 16 | 25Sep2015 | 8 | 24.31 | 29.42 |
| | | Bugang (WS1667) | 17 | 27Aug2015 | 10 | 26.33 | 23.98 |
| | | Sacheon (WS1669) | 19 | 26Aug2015 | 8 | 29.78 | 25.49 |
| | | Shanghai (WS1665) | 15 | 28Aug2015 | 9 | 25.25 | 25.68 |
| | | Total | | | 341 | 27.78 | 28.58 |

**Table S5.** Parameter estimates for fixed and random effects from Bayesian GLMM models of gut length in seven subgroups, with individual ID included as a random effect to account for unmeasured individual variation. Effect sizes of each subgroup (SG indicates the same as "genetic background" in subsequent analyses) and male are given relative to the model intercepts (NJPN1 and female, respectively) and are the posterior mean of 100,000 sampled iterations. The lower and upper CIs are given at the 95% significance level. Four MCMC chains were run for 105,000 samples, with a 5,000 sample burn-in, followed by every 4th sample being used. This gave a total of 100,000 samples used.

| Fixed effects | Estimate | Est. Error | l-95% CI | u-95% CI | R-hat |
|---|---|---|---|---|---|
| **Intercept** | **2.71** | **0.17** | **2.37** | **3.05** | **1.00** |
| **Sex: male** | **-0.11** | **0.03** | **-0.16** | **-0.06** | **1.00** |
| **Standard length** | **0.03** | **0.01** | **0.02** | **0.04** | **1.00** |
| **SG - NJPN2** | **-0.28** | **0.05** | **-0.38** | **-0.17** | **1.00** |
| **- SJPN1** | **-0.11** | **0.04** | **-0.20** | **-0.02** | **1.00** |
| **- SJPN2** | **-0.13** | **0.04** | **-0.22** | **-0.05** | **1.00** |
| **- SJPN3** | **-0.23** | **0.05** | **-0.32** | **-0.13** | **1.00** |
| **- SJPN4** | **-0.26** | **0.04** | **-0.34** | **-0.19** | **1.00** |
| **- KOR** | **-0.25** | **0.05** | **-0.35** | **-0.16** | **1.00** |

| Random effect | Estimate | Est. Error | l-95% CI | u-95% CI | R-hat |
|---|---|---|---|---|---|
| **ID** | **0.18** | **0.04** | **0.09** | **0.23** | **1.00** |

**Table S6**. Parameter estimates for fixed and random effects from Bayesian GLMM model of gut lengths in seasons and genetic backgrounds, with individual ID and year (2015 or 2021) included as random effects to account for unmeasured individual variation and the differences between years, respectively. Effect sizes of winter, genetic backgrounds (NJPN2 and SJPN) and male are given relative to the model intercepts (summer, NJPN1 and female, respectively) and are the posterior mean of 10,000 sampled iterations. The lower and upper CIs are given at the 95% significance level. Four MCMC chains were run for 11,000 samples, with a 1,000 sample burn-in, followed by every 4th sample being used. This gave a total of 10,000 samples used.

Response variable: Gut length

| Fixed effects | Estimate | Est. Error | l-95% CI | u-95% CI | R-hat |
|---|---|---|---|---|---|
| **Intercept** | **2.54** | **0.77** | **1.36** | **4.12** | **1.01** |
| **Sex: male** | **-0.08** | **0.03** | **-0.13** | **-0.02** | **1.00** |
| **Standard length** | **0.04** | **0.01** | **0.02** | **0.05** | **1.00** |
| Season: winter | 0.03 | 0.05 | -0.06 | 0.12 | 1.00 |
| **Genetic background: NJPN2** | **-0.20** | **0.04** | **-0.27** | **-0.12** | **1.00** |
| **SJPN** | **-0.14** | **0.04** | **-0.21** | **-0.06** | **1.00** |
| Interaction: winter × NJPN2 | -0.09 | 0.06 | -0.22 | 0.03 | 1.00 |
| **winter × SJPN** | **-0.19** | **0.06** | **-0.32** | **-0.07** | **1.00** |

| Random effects | Estimate | Est. Error | l-95% CI | u-95% CI | R-hat |
|---|---|---|---|---|---|
| **ID** | **0.11** | **0.05** | **0.01** | **0.18** | **1.00** |
| **Year** | **0.62** | **0.98** | **0.01** | **3.47** | **1.00** |

This model includes the interaction between season and genetic background.

**Table S7**. List of genes in the vicinity of seasonally methylated regions. The gene symbols in bold were supported by both Reactome and Gene ontology (GO) analyses. The asterisks and dagger mean that genes with DMRs in the vicinity were the same and the opposite results in WGBS, respectively.

Methylated regions in summer within 2 kbp from gene

| | Peak (bp) | | | | Position (bp) | | | Methylated region from 5' or 3' UTRs | |
|---|---|---|---|---|---|---|---|---|---|
| Chr. | Start | End | Length | Symbol | Start | End | Strand | Location | Distance (bp) |
| chr1 | 2224850 | 2225050 | 200 | armc3 | 2219570 | 2227531 | reverse | body | 2481 |
| chr1 | 2635750 | 2635900 | 150 | EVI5L | 2635046 | 2646741 | forward | body | 704 |
| chr1 | 11574850 | 11575050 | 200 | tyrp1b* | 11575165 | 11583484 | forward | upstream | -315 |
| chr1 | 25655450 | 25655650 | 200 | cacna1i | 25647228 | 25664946 | forward | body | 8222 |
| chr2 | 206450 | 206650 | 200 | DMD | 204721 | 208200 | forward | body | 1729 |
| chr2 | 10909450 | 10909600 | 150 | agap1 | 10900109 | 10911780 | forward | body | 9341 |
| chr3 | 32933250 | 32933400 | 150 | abcc12 | 32920227 | 32941583 | reverse | body | 8183 |
| chr4 | 18235550 | 18235800 | 250 | stxbp3* | 18231997 | 18240790 | forward | body | 3553 |
| chr4 | 27005500 | 27005650 | 150 | tjp3 | 27001627 | 27018892 | reverse | body | 13242 |
| chr5 | 1831650 | 1831900 | 250 | SZRD1 | 1822897 | 1831277 | forward | downstream | -373 |
| chr5 | 23619100 | 23619300 | 200 | cdc42 | 23615317 | 23622399 | reverse | body | 3099 |
| chr6 | 510150 | 510300 | 150 | ppfibp1b* | 506802 | 528444 | forward | body | 3348 |
| chr7 | 8218100 | 8218300 | 200 | PRDM16 | 8205031 | 8252270 | reverse | body | 33970 |
| **chr7** | **30322050** | **30322350** | **300** | **plxnb3*** | **30323235** | **30355796** | **forward** | **upstream** | **-1185** |
| **chr7** | **31962950** | **31963100** | **150** | **plxna3** | **31962373** | **31977272** | **reverse** | **body** | **14172** |
| chr8 | 8598300 | 8598450 | 150 | col5a3a | 8598717 | 8626264 | reverse | downstream | -267 |
| chr9 | 11811250 | 11811450 | 200 | gna14a | 11805116 | 11815981 | forward | body | 6134 |
| chr9 | 14094050 | 14094200 | 150 | iqgap2* | 14078174 | 14100828 | forward | body | 15876 |
| chr10 | 1453200 | 1453400 | 200 | fam198b* | 1450732 | 1453540 | forward | body | 2468 |
| chr10 | 3166250 | 3166600 | 350 | exosc3* | 3164842 | 3169191 | forward | body | 1408 |
| chr10 | 13738600 | 13738750 | 150 | eef1g | 13730304 | 13740531 | reverse | body | 1781 |
| chr10 | 13738600 | 13738750 | 150 | si:ch211-175m2.5 | 13740401 | 13745216 | forward | upstream | -1801 |
| chr11 | 3549900 | 3550050 | 150 | GRIK5 | 3539189 | 3553965 | forward | body | 10711 |
| chr11 | 16935150 | 16935300 | 150 | macf1a* | 16889162 | 16935571 | reverse | body | 271 |
| chr11 | 19898200 | 19898350 | 150 | meaf6 | 19898200 | 19908217 | forward | body | 0 |
| chr12 | 2586350 | 2586500 | 150 | SBNO1* | 2584185 | 2597462 | forward | body | 2165 |
| chr12 | 6050700 | 6050850 | 150 | col5a1 | 6019455 | 6062286 | forward | body | 31245 |

| | | | | | | | | | |
|---|---|---|---|---|---|---|---|---|---|
| chr13 | 5057000 | 5057200 | 200 | ehd2b | 5050697 | 5058169 | forward | body | 6303 |
| chr13 | 11409600 | 11409800 | 200 | dixdc1a | 11405249 | 11415353 | forward | body | 4351 |
| chr13 | 17220250 | 17220450 | 200 | PDE3A | 17222255 | 17223717 | forward | upstream | -2005 |
| chr13 | 22390050 | 22390250 | 200 | hip1rb* | 22380619 | 22394235 | reverse | body | 3985 |
| chr13 | 28193100 | 28193250 | 150 | SAMSN1* | 28185234 | 28203455 | forward | body | 7866 |
| chr14 | 9469650 | 9469850 | 200 | waif2 | 9464834 | 9488924 | forward | body | 4816 |
| chr14 | 9950000 | 9950150 | 150 | kl | 9951318 | 9960110 | forward | upstream | -1318 |
| chr14 | 9950000 | 9950150 | 150 | pds5b | 9929947 | 9951125 | forward | body | 20053 |
| chr14 | 21651450 | 21651650 | 200 | cenpv* | 21653018 | 21655297 | reverse | downstream | -1368 |
| chr14 | 24831250 | 24831400 | 150 | ppm1e | 24823963 | 24833149 | forward | body | 7287 |
| chr15 | 18460350 | 18460550 | 200 | ank3a | 18428781 | 18482128 | reverse | body | 21578 |
| chr16 | 5536900 | 5537050 | 150 | ptpn2a | 5528464 | 5535099 | forward | downstream | -1801 |
| chr16 | 5536900 | 5537050 | 150 | cep76 | 5538011 | 5546259 | forward | upstream | -1111 |
| chr16 | 10376700 | 10376950 | 250 | CLPTM1L | 10373182 | 10380746 | forward | body | 3518 |
| chr16 | 12724900 | 12725050 | 150 | adcy2b | 12718718 | 12730302 | forward | body | 6182 |
| chr17 | 12386800 | 12387100 | 300 | aktip | 12386754 | 12387101 | reverse | body | 1 |
| chr17 | 12386800 | 12387100 | 300 | aktip | 12387160 | 12388209 | reverse | downstream | -60 |
| chr17 | 22023450 | 22023700 | 250 | pip5k1cb* | 22015212 | 22029750 | reverse | body | 6050 |
| chr19 | 13149300 | 13149550 | 250 | eef2k* | 13147935 | 13159456 | reverse | body | 9906 |
| chr20 | 1474700 | 1474900 | 200 | cubn | 1456113 | 1511393 | reverse | body | 36493 |
| chr20 | 8293200 | 8293350 | 150 | itgb1a* | 8291957 | 8306080 | reverse | body | 12730 |
| chr20 | 11838200 | 11838350 | 150 | nsmaf | 11824680 | 11838960 | forward | body | 13520 |
| chr20 | 11838200 | 11838350 | 150 | cngb3 | 11839417 | 11849956 | reverse | downstream | -1067 |
| chr20 | 13893200 | 13893400 | 200 | zfhx4 | 13888533 | 13899412 | forward | body | 4667 |
| chr20 | 19030250 | 19030450 | 200 | chst2b | 19026878 | 19028275 | reverse | upstream | -1975 |
| chr20 | 19030250 | 19030450 | 200 | trpc1 | 19032433 | 19050511 | forward | upstream | -2183 |
| chr20 | 23592650 | 23592800 | 150 | nceh1b*† | 23593106 | 23599431 | reverse | downstream | -306 |
| chr21 | 4195250 | 4195650 | 400 | MAP4K4* | 4195823 | 4196364 | reverse | downstream | -173 |
| chr21 | 6690950 | 6691150 | 200 | cwc22 | 6690772 | 6706864 | reverse | body | 15714 |
| chr21 | 21724000 | 21724200 | 200 | ppig* | 21715197 | 21722782 | forward | downstream | -1218 |
| chr21 | 23972950 | 23973100 | 150 | egfl6 | 23966397 | 23979776 | forward | body | 6553 |
| chr22 | 7905100 | 7905300 | 200 | lclat1* | 7903821 | 7915438 | forward | body | 1279 |
| chr22 | 16162150 | 16162350 | 200 | SCAF8* | 16161695 | 16174043 | reverse | body | 11693 |
| **chr23** | **14882850** | **14883100** | **250** | **plxna4** | **14881290** | **14885450** | **reverse** | **body** | **2350** |

| | | | | | | | | | |
|---|---|---|---|---|---|---|---|---|---|
| chr23 | 22623800 | 22623950 | 150 | TRHDE | 22620605 | 22625230 | reverse | body | 1280 |
| chr24 | 1577850 | 1578000 | 150 | hey1* | 1576857 | 1579313 | reverse | body | 1313 |
| chr24 | 4634100 | 4634350 | 250 | rgs6 | 4629093 | 4647171 | forward | body | 5007 |
| chr24 | 9254650 | 9254850 | 200 | prox1a* | 9251700 | 9255162 | reverse | body | 312 |
| chr24 | 10164950 | 10165100 | 150 | prpf39 | 10164324 | 10178663 | reverse | body | 13563 |
| chr24 | 13505200 | 13505400 | 200 | pnocb | 13507094 | 13512960 | reverse | downstream | -1694 |
| chr24 | 13505200 | 13505400 | 200 | znf395b | 13493521 | 13505414 | forward | body | 11679 |
| chr24 | 21059200 | 21059500 | 300 | epha7 | 21055708 | 21075901 | reverse | body | 16401 |

### Methylated regions in the winter within 2 kbp from the gene

| | Peak (bp) | | | | Position (bp) | | | Methylated region from 5' or 3' UTRs | |
|---|---|---|---|---|---|---|---|---|---|
| Chr. | Start | End | Length | Symbol | Start | End | Strand | Location | Distance (bp) |
| chr2 | 9495700 | 9496050 | 350 | RNH1 | 9491265 | 9501818 | reverse | body | 5768 |
| chr11 | 7730400 | 7730600 | 200 | CSMD3 | 7730225 | 7744745 | reverse | body | 14145 |

**Table S8.** Top 10 terms in the Reactome and GO (biological process) estimated using Enrichr (*42, 43*) analysis for genes in the vicinity of winter-methylated regions. Overlap shows a proportion of input set of genes in annotated gene sets. *P*-value and adjusted *P*-value are calculated using Fisher's exact test and the Benjamini-Hochberg method for correction for multiple hypotheses testing; adjusted *P*-values <0.05 are shown in bold. Enrichr was performed on 27 Jan. 2019.

| Reactome term | Overlap | *P*-value | Adjusted *P*-value |
|---|---|---|---|
| **Axon guidance_Homo sapiens_R-HSA-422475** | **11/515** | **1.04E-06** | **1.69E-04** |
| **Semaphorin interactions_Homo sapiens_R-HSA-373755** | **6/67** | **9.56E-08** | **3.13E-05** |
| **Developmental biology_Homo sapiens_R-HSA-1266738** | **11/786** | **5.54E-05** | **3.62E-03** |
| **Other semaphorin interactions_Homo sapiens_R-HSA-416700** | **3/19** | **3.35E-05** | **2.74E-03** |
| **SEMA3A-Plexin repulsion signaling by inhibiting Integrin adhesion_Homo sapiens_R-HSA-399955** | **3/14** | **1.27E-05** | **1.39E-03** |
| **RHO GTPases activate IQGAPs_Homo sapiens_R-HSA-5626467** | **2/11** | **5.96E-04** | **3.25E-02** |
| Sema3A PAK dependent Axon repulsion_Homo sapiens_R-HSA-399954 | 2/16 | 1.29E-03 | 5.26E-02 |
| CRMPs in Sema3A signaling_Homo sapiens_R-HSA-399956 | 2/16 | 1.29E-03 | 5.26E-02 |
| Hemostasis_Homo sapiens_R-HSA-109582 | 6/552 | 1.04E-02 | 2.63E-01 |
| Non-integrin membrane-ECM interactions_Homo sapiens_R-HSA-3000171 | 2/42 | 8.73E-03 | 2.63E-01 |

| Gene ontology term in the biological process | Overlap | *P*-value | Adjusted *P*-value |
|---|---|---|---|
| **Semaphorin-plexin signaling pathway involved in axon guidance (GO:1902287)** | **3/10** | **4.24E-06** | **3.02E-03** |
| **Semaphorin-plexin signaling pathway involved in neuron projection guidance (GO:1902285)** | **3/14** | **1.27E-05** | **3.30E-03** |
| **Semaphorin-plexin signaling pathway (GO:0071526)** | **3/30** | **1.37E-04** | **1.77E-02** |
| **Positive regulation of sodium ion transmembrane transport (GO:1902307)** | **2/7** | **2.30E-04** | **2.22E-02** |
| **Motor neuron axon guidance (GO:0008045)** | **2/11** | **5.96E-04** | **3.56E-02** |
| **Positive regulation of sodium ion transmembrane transporter activity (GO:2000651)** | **2/12** | **7.14E-04** | **3.96E-02** |
| **Positive regulation of axonogenesis (GO:0050772)** | **4/38** | **7.79E-06** | **3.02E-03** |
| **Regulation of platelet-derived growth factor receptor-beta signaling pathway (GO:2000586)** | **2/10** | **4.89E-04** | **3.29E-02** |
| **Regulation of axon guidance (GO:1902667)** | **2/14** | **9.80E-04** | **4.40E-02** |
| **Plasma membrane organization (GO:0007009)** | **3/37** | **2.57E-04** | **2.22E-02** |

**Table S9**. Parameter estimates from the Bayesian GLM models of standard and gut lengths in offspring generated by heterozygous mating of CRISPR-induced mutant medaka. Effect sizes of each genotype and male are given relative to the model intercepts (wild type [+/+] and female, respectively) and are the posterior mean of 4,000 sampled iterations. The lower and upper CIs are given at the 95% significance level. Four MCMC chains were run for 2,000 samples, with a 1,000 sample burn-in, followed by every sample being used. This gave a total of 4,000 samples used.

Response variable: Standard length

| Variable | Estimate | Est. Error | l-95% CI | u-95% CI | R-hat |
|---|---|---|---|---|---|
| **Intercept** | **3.13** | **0.11** | **2.9** | **3.36** | **1.00** |
| Sex: male | -0.02 | 0.02 | -0.07 | 0.02 | 1.01 |
| Day after hatching | 0 | 0 | 0 | 0 | 1.00 |
| +/- | -0.03 | 0.02 | -0.07 | 0.02 | 1.00 |
| **-/-** | **-0.07** | **0.03** | **-0.12** | **-0.02** | **1.00** |

Response variable: Gut length

| Variable | Estimate | Est. Error | l-95% CI | u-95% CI | R-hat |
|---|---|---|---|---|---|
| **Intercept** | **2.16** | **0.58** | **0.99** | **3.32** | **1.00** |
| **Sex: male** | **-0.15** | **0.05** | **-0.25** | **-0.05** | **1.00** |
| Day after hatch | 0 | 0 | 0 | 0.01 | 1.00 |
| Standard length | 0.02 | 0.02 | -0.03 | 0.07 | 1.00 |
| +/- | -1.56 | 0.8 | -3.08 | 0.02 | 1.00 |
| -/- | -1.92 | 1.05 | -3.95 | 0.14 | 1.00 |
| SL × +/- | 0.07 | 0.04 | 0 | 0.14 | 1.00 |
| SL × -/- | 0.09 | 0.05 | -0.01 | 0.19 | 1.00 |

This model includes the interaction between standard length and genotype.

Response variable: Gut length

| Variable | Estimate | Est. Error | l-95% CI | u-95% CI | R-hat |
|---|---|---|---|---|---|
| **Intercept** | **2.17** | **0.56** | **1.08** | **3.28** | **1.00** |
| **Sex: male** | **-0.15** | **0.05** | **-0.25** | **-0.06** | **1.00** |
| Day after hatch | 0 | 0 | 0 | 0.01 | 1.00 |
| Standard length | 0.02 | 0.02 | -0.03 | 0.06 | 1.00 |
| **+/-, -/-*** | **-1.67** | **0.67** | **-3.00** | **-0.31** | **1.00** |
| **SL × ( +/-, -/-)** | **0.08** | **0.03** | **0.01** | **0.14** | **1.00** |

*This model hypothesizes the effect of mutation as a dominance.

**Table S10**. Parameter estimates for fixed and random effects from Bayesian GLMM model of gut length in *Ppp3r1* genotypes, with individual ID and genetic background included as random effects to account for unmeasured individual variation and population stratification, respectively. Effect sizes of each genotype and male are given relative to the model intercepts (C/C and female, respectively) and are the posterior mean of 100,000 sampled iterations. The lower and upper CIs are given at the 95% significance level. Four MCMC chains were run for 105,000 samples, with a 5,000 sample burn-in, followed by every 4th sample being used. This gave a total of 100,000 samples used.

| Fixed effects | Estimate | Est. Error | l-95% CI | u-95% CI | R-hat |
|---|---|---|---|---|---|
| **Intercept** | **2.24** | **0.20** | **1.86** | **2.63** | **1.00** |
| **Sex: male** | **-0.10** | **0.02** | **-0.14** | **-0.05** | **1.00** |
| **Standard length** | **0.04** | **0.01** | **0.03** | **0.05** | **1.00** |
| *Ppp3r1* - C/T | 0.15 | 0.14 | -0.12 | 0.42 | 1.00 |
| **- T/T** | **0.23** | **0.06** | **0.11** | **0.34** | **1.00** |

| Random effects | Estimate | Est. Error | l-95% CI | u-95% CI | R-hat |
|---|---|---|---|---|---|
| **ID** | **0.14** | **0.04** | **0.03** | **0.19** | **1.00** |
| **Genetic background** | **0.13** | **0.02** | **0.09** | **0.17** | **1.00** |

**Table S11**. Parameter estimates for fixed and random effects from Bayesian GLMM models of gut length in *Ppp3r1* expression, with individual ID and genetic background included as random effects to account for unmeasured individual variation and population stratification, respectively. The effect size of male is given relative to the model intercept (female). The lower and upper CIs are given at the 95% significance level. Four MCMC chains were run for 105,000 samples, with a 5,000 sample burn-in, followed by every 4th sample being used. This gave a total of 100,000 samples used.

Model 1 (Not controlled for genetic background)

| Fixed effects | Estimate | Est. Error | l-95% CI | u-95% CI | R-hat |
|---|---|---|---|---|---|
| Intercept | 1.79 | 0.92 | -0.05 | 3.61 | 1.00 |
| Sex: male | -0.18 | 0.15 | -0.47 | 0.12 | 1.00 |
| Standard length | 0.06 | 0.04 | -0.01 | 0.13 | 1.00 |
| **Relative *Ppp3r1* expression** | **0.42** | **0.12** | **0.19** | **0.66** | **1.00** |

| Random effect | Estimate | Est. Error | l-95% CI | u-95% CI | R-hat |
|---|---|---|---|---|---|
| **ID** | **0.15** | **0.08** | **0.01** | **0.33** | **1.00** |

Model 2 (Controlled for genetic background)

| Fixed effects | Estimate | Est. Error | l-95% CI | u-95% CI | R-hat |
|---|---|---|---|---|---|
| Intercept | 1.76 | 1.17 | -0.63 | 4.07 | 1.00 |
| Sex: male | -0.13 | 0.16 | -0.45 | 0.19 | 1.00 |
| Standard length | 0.06 | 0.04 | -0.02 | 0.14 | 1.00 |
| Relative *Ppp3r1* expression | 0.29 | 0.22 | -0.18 | 0.68 | 1.00 |

| Random effects | Estimate | Est. Error | l-95% CI | u-95% CI | R-hat |
|---|---|---|---|---|---|
| **ID** | **0.14** | **0.08** | **0.01** | **0.32** | **1.00** |
| **Genetic background** | **0.46** | **0.54** | **0.01** | **2.00** | **1.00** |

**Table S12.** List of the top 10 SNPs associated with the gut length variation.

| Chr | Pos. | Weir and Cockerham's $F_{ST}$ | Associated *P*-value† | Corrected *P*-value‡ | Gene Symbol | Ensembl Gene ID |
|---|---|---|---|---|---|---|
| 1 | 635,654 | 0.892 | 4.23E-13 | 7.00E-07 | *Ppp3r1* | ENSORLG00000025788* |
| 16 | 27,832,716 | 0.510 | 1.96E-11 | 8.10E-06 | *si:ch73-95l15.5* | ENSORLG00000030239 |
| 3 | 37,278,080 | 0.691 | 2.51E-11 | 9.50E-06 | *Spg21* | ENSORLG00000018308 |
| 6 | 24,039,480 | 0.965 | 4.45E-11 | 1.52E-05 | *Slc7a5* | ENSORLG00000012624 |
| 21 | 3,752,404 | 0.568 | 6.72E-11 | 1.98E-05 | *Asb11* | ENSORLG00000008942 |
| 19 | 134,355 | 0.903 | 1.13E-10 | 2.67E-05 | – | – |
| 8 | 18,083,252 | 0.926 | 1.39E-10 | 3.10E-05 | *Vill* | ENSORLG00000009593 |
| 16 | 27,832,682 | 0.226 | 1.40E-10 | 3.11E-05 | *si:ch73-95l15.5* | ENSORLG00000030239 |
| 22 | 1,476,147 | 0.780 | 3.17E-10 | 5.39E-05 | *Pacs2* | ENSORLG00000009340 |
| 12 | 2,567,193 | 0.811 | 1.51E-09 | 1.54E-04 | *Rilpl1* | ENSORLG00000001553 |

† Uncorrected *P*-values obtained from association analysis.
‡ Corrected *P*-values calculated by permutation tests with resampling 10,000,000 times.
* This gene is paralogous to the *Ppp3r1a* gene on chromosome 15.

**Table S13.** Parameter estimates for fixed and random effects from Bayesian GLMM models of gut length in the number of CpG sites and *Ppp3r1* genotypes, with genetic background included as a random effect to account for population stratification. Effect size of *Ppp3r1*-T/T is given relative to the model intercept (*Ppp3r1*-C/C). The lower and upper CIs are given at the 95% significance level. Four MCMC chains were run for 105,000 samples, with a 5,000 sample burn-in, followed by every 4th sample being used. This gave a total of 100,000 samples used.

| Fixed effects | Estimate | Est. Error | l-95% CI | u-95% CI | R-hat |
|---|---|---|---|---|---|
| **Intercept** | **2.86** | **0.19** | **2.46** | **3.21** | **1.00** |
| **Num. CpG** | **0.06** | **0.02** | **0.02** | **0.11** | **1.00** |
| ***Ppp3r1*-T/T** | **0.32** | **0.09** | **0.15** | **0.49** | **1.00** |

| Random effect | Estimate | Est. Error | l-95% CI | u-95% CI | R-hat |
|---|---|---|---|---|---|
| **Genetic background** | **0.05** | **0.04** | **0.00** | **0.16** | 1.00 |
